# Synovial 1 Sarcoma X Breakpoint Protein Uses a Cryptic Groove to Selectively Recognize H2AK119Ub Nucleosomes

**DOI:** 10.1101/2023.07.08.548231

**Authors:** Zebin Tong, Huasong Ai, Ziyu Xu, Kezhang He, Guo-Chao Chu, Qiang Shi, Zhiheng Deng, Qiaomei Xue, Maoshen Sun, Yunxiang Du, Lujun Liang, Jia-Bin Li, Man Pan, Lei Liu

## Abstract

The cancer-specific fusion oncoprotein SS18-SSX1 disturbs chromatin accessibility by hijacking the BAF complex from the promoters and enhancers to the polycomb-repressed chromatin regions. This process relies on the selective recognition of H2AK119Ub nucleosomes by SSX1. However, the mechanism by which SSX1 selective recognition of H2AK119Ub nucleosomes in the absence of ubiquitin (Ub)-binding capacity remains unknown. Here we report the cryo-EM structure of SSX1 bound to H2AK119Ub nucleosomes at 3.1 Å resolution. Combined *in vitro* biochemical and cellular assays revealed that the Ub recognition by SSX1 is unique and depends on a cryptic basic groove formed by H3 and the Ub motif on the H2AK119 site. Moreover, this unorthodox binding mode of SSX1 induces DNA unwrapping at the entry/exit sites. Together, our results describe a unique mode of site-specific ubiquitinated nucleosome recognition that underlies the specific hijacking of the BAF complex to polycomb regions by SS18-SSX1 in synovial sarcoma.

## Introduction

The homeostatic maintenance of chromatin accessibility, a process important to gene expression and cellular identity regulation, is exquisitely controlled by multiple chromatin-binding factors such as chromatin remodelers and histone-modifying enzymes^1^. The competition of different chromatin-binding factors for nucleosomes has established a dynamic balance of accessible and inaccessible chromatin processes^1^. A representative example involves the opposition between ATP-dependent chromatin-remodeling BAF complexes and polycomb repression complexes (PRCs), which help to establish accessible and inaccessible chromatin regions, respectively^2^. BAF complexes remodel nucleosomes, enhancing DNA accessibility at the promoters and enhancers^3,4^. In contrast, PRC1 catalyses the monoubiquitination of H2A at lysine 119 (H2AK119Ub)^5^, which recruits and activates PRC2 to drive the trimethylation of histone H3 at lysine 27 (H3K27me3)^6,7^, leading to polycomb-compacted and inaccessible chromatin^8,9^. Mechanistic studies have provided molecular insights into how PRC2 and its auxiliary subunits, JARID2 and AEBP2, selectively target H2AK119Ub nucleosomes^10^. Two major interactions are revealed: the anchoring of an arginine- or lysine-rich motif to the nucleosomal H2A/H2B acidic patch and the interaction of the canonical ubiquitin-binding domain (e.g., Ub-interacting motif (UIM) domain in JARID2, zinc finger in AEBP2) with the Ub on histone. This divalent recognition mode is also shared with other ubiquitinated nucleosome-recruiting effector proteins, such as 53BP1^11^ and BARD1^12,13^, which recognize H2AK15Ub nucleosomes to regulate the DNA double-strand breaks (DSBs) response pathway.

The disruption of chromatin-binding factors that regulate chromatin accessibility often accompanies diseases such as cancer^2,14^. An important example is the dysregulation of the BAF complex by the fusion oncoprotein SS18-SSX1, which is produced by the genetic fusion of SS18 (one of the subunits of the BAF complex, residues 1-379) and the gene silencing factor synovial sarcoma X breakpoint 1 (SSX1, residues 111-188) due to pathognomonic t(X;18)(p11.2;q11.2) chromosomal translocation in synovial sarcoma^15,16^. Nearly 100% of human synovial sarcoma cases are caused by the SS18-SSX1 fusion oncoprotein. The incorporation of SS18-SSX into BAF complexes (known as ssBAF) hijacks BAF complexes from chromatin promoters or enhancers to H2AK119Ub enriched polycomb-repressed regions, which mediates the eviction of PRCs from chromatin and activates polycomb-repressed genes (e.g., the activation of the oncogene *SOX2*)^17–19^. An important recent discovery is that mistargeting of ssBAF is achieved through the selective recognition of H2AK119Ub nucleosomes by SSX1 residues 111-188 in the fusion oncoprotein SS18-SSX1^20^. However, the mechanism of SSX1 recruitment through the selective recognition of H2AK119Ub nucleosomes remains unknown. A particular puzzle is that SSX1 neither binds to free Ub *in vitro*^20^, nor has any currently-known Ub-binding domain (e.g., UIM, zinc finger, Ub-associated domain (UBA)) according to bioinformatic predictions, implying the existence of a unique mode of site-specific ubiquitinated nucleosome recognition.

Here, we determined the cryo-EM structure of SSX1 bound to H2AK119Ub nucleosomes at a resolution of 3.1 Å. This structure reveals three interaction regions between SSX1 and H2AK119Ub nucleosomes. First, SSX1 N-terminal basic regions (residues W164, R167, L168, R169) are anchored into the H2A/H2B acidic patch. Second, the aromatic amino acid (residue Y177) in the middle of SSX1 interacts with the hydrophobic pocket formed by H2A α2 and α3 helixes. Third, the SSX1 C-terminal acidic region binds to a cryptic basic groove formed by H3 (residues R49, R52, K56) and the Ub motif (residues R42, R72, R74) on H2AK119. This H2AK119Ub nucleosome-specific “H3-Ub^H2AK119^” basic groove is the structural basis by which SSX1 selectively recognizes H2AK119Ub nucleosomes in the absence of any Ub-binding domains. Moreover, the binding of SSX1 to the H3-Ub^H2AK119^ basic groove is observed to induce nucleosomal DNA unwrapping. Collectively, our structural and biochemical data reveal a unique recognition model for SSX1 recruitment to H2AK119Ub nucleosomes that underlies the specific hijacking of the BAF complex to polycomb regions by SS18-SSX1 in synovial sarcoma.

## Results

### SSX1 preferentially binds to H2AK119Ub nucleosomes

Recent studies have shown that SSX1 has a higher affinity for H2AK119Ub nucleosomes than for H2BK120Ub nucleosomes^20^. It remains unknown whether SSX1 can recognize other ubiquitinated nucleosomes, such as H2AK127Ub and H3K14Ub whose Ub is located near the nucleosomal DNA entry/exit site, or H2AK15Ub whose Ub is located on the opposite side of the nucleosomal DNA entry/exit sites (**Fig. 1a**). In this context, we expressed SSX1(111-188) which is a segment of SSX1 protein fused to SS18 and sufficient for selective recognition of H2AK119Ub nucleosomes. Moreover, the GST tag was fused at SSX1(111-188) N-terminus for pull-down experiments (**Extended Data Fig. 1a**). The purified GST-SSX1 protein was used to pull down five chemically ubiquitinated nucleosomes (**Supplementary Fig. 17c.**)^21,22^. As shown in **Fig. 1b**, GST-SSX1 successfully pulled down histone bands for H2AK119Ub nucleosomes but not for H2BK120Ub nucleosomes, consistent with previous AlphaLISA results^20^. Moreover, GST-SSX1 showed no pull-down histone bands for H2AK15Ub, H2AK127Ub, and H3K14Ub nucleosomes, indicating that SSX1 specifically binds ubiquitinated nucleosomes at H2AK119 over all the other sites. Notably, H2AK127 differs from H2AK119 by only seven interval residues (**Fig. 1b**), suggesting that the specific recognition of H2AK119Ub nucleosomes by SSX1 is not only regioselective at the nucleosomal DNA entry/exit sites, but also strictly site-selective at H2AK119 only.

**Fig. 1.**
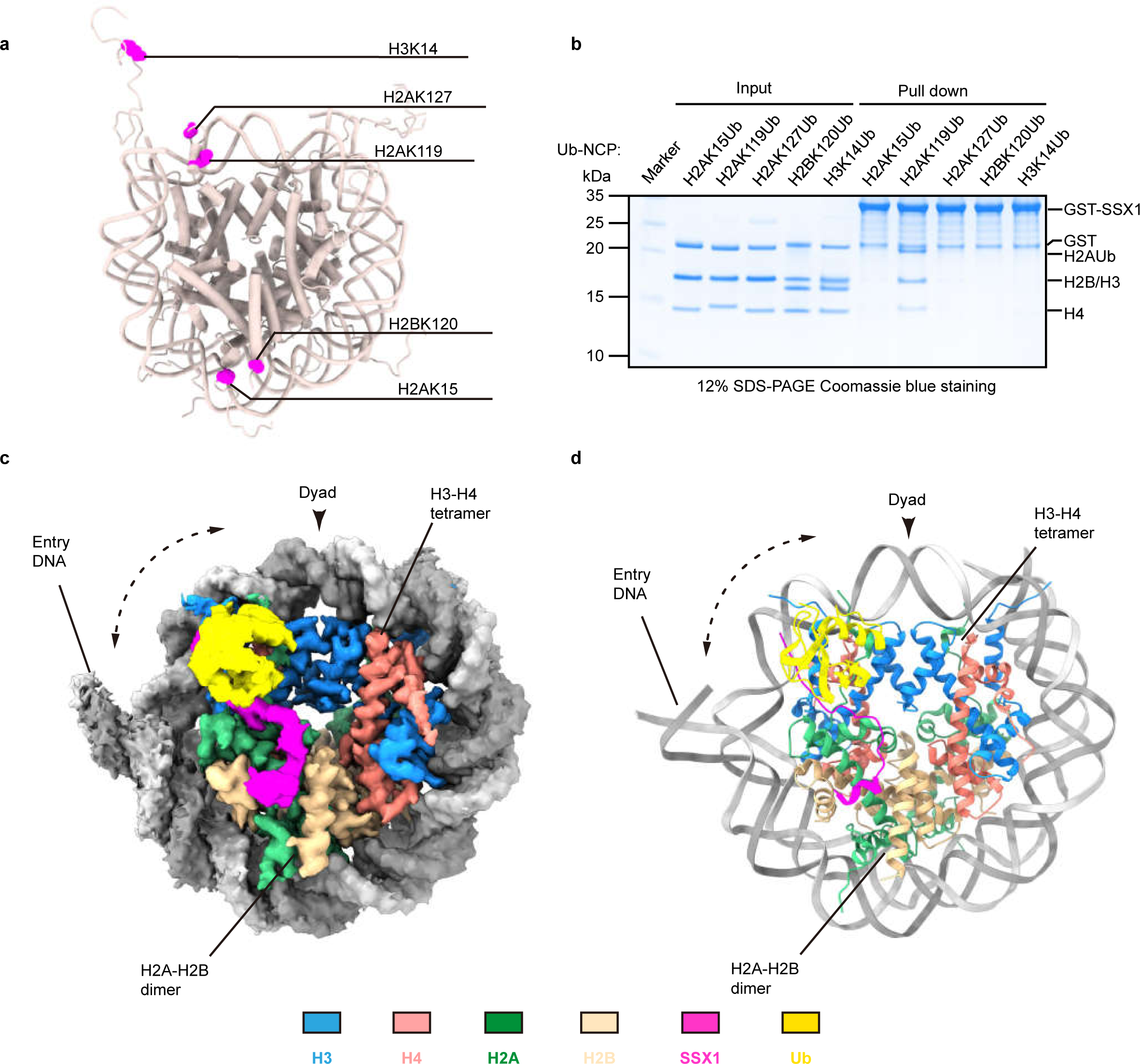
Cryo-EM structure of SSX1 bound to H2AK119Ub nucleosomes. **a.** Schematic representation of the five ubiquitinated sites on the nucleosomes, including H2AK15, H2AK119, H2AK127, H2BK120, and H3K14. **b.** Pull-down experiments of GST-SSX1 incubated with five ubiquitinated nucleosomes (As shown in **a**) and visualized with Coomassie blue-staining. These gels are representative of five independent experiments. The gel source data is shown in Source Data Fig.1. **c.** The 3.1 Å Cryo-EM density map of SSX1-H2AK119Ub nucleosome complex. **d.** Atomic model of the SSX1-H2AK119Ub nucleosome complex. The structure is depicted in cartoon with round helices. The SSX1 was coloured in magenta, ubiquitin in yellow, DNA double helix in light grey and dark grey, respectively, histone H2A in media sea green, histone H2B in burly wood, histone H3 in dodger blue, histone H4 in salmon.

### Structural overview of the SSX1-H2AK119Ub nucleosome complex

To elucidate the molecular mechanism by which SSX1 selectively binds to H2AK119Ub nucleosomes, GST-SSX1 was incubated with H2AK119Ub nucleosomes and the complex was subjected to cryo-EM single-particle analysis **(Extended Data Fig. 1b, c, f)**. A total of 4805 stacks were collected on a 300 kV Titan Krios microscope and processed with RELION 3.1 software^23^ (**Extended Data Fig. 2a-c)**. By carefully processing the cryo-EM data of SSX1-H2AK119Ub nucleosome complex, we obtained two types of SSX1-H2AK119Ub nucleosome complex: the first is the SSX1:H2AK119Ub NCP=2:1 (hereafter 2:1 complex) bound conformation at a resolution of 3.6 Å **(Extended Data Fig. 3c)**, utilizing 60624 particles. In this map, it is evident that two SSX1 molecules bind to both sides of the nucleosome discs and the densities of Ub on both sides of the nucleosome are visible **(Extended Data Fig. 3b)**. The second was the SSX1:H2AK119Ub NCP=1:1 bound conformation (hereafter 1:1 complex) at a resolution of 3.1 Å reconstructed with 195,091 particles, which is also the main conformation **(Extended Data Fig. 3a, c, d)**. In this map, the nucleosome in the complex forms a symmetrical coin shape, and the densities of the secondary structures of the four core histones can be clearly identified **(Extended Data Fig. 3e)**. The extra density on one side of the nucleosome can be attributed to Ub and SSX1 **(Fig. 1c)**. The density of the GST moiety is too low to visualize despite the blurred density outside the nucleosome disc in the 2D and 3D classifications **(Extended Data Fig. 1d, e)**. The final reconstruction showed sufficiently detailed and well-resolved residue density to allow us to build a structural model of the SSX1-H2AK119Ub nucleosome complex, including H2A(13-119), H2B(34-142), H3(38-134), H4(19-101), Ub(1-74) and SSX1(162-184) **(Fig. 1d and Extended Data Fig. 3e)**. In this model, SSX1(162-184) binds to nucleosomes in a snake shape, travelling approximately 60 Å from the nucleosomal H2A/H2B acidic patch to the histone H3 αN-helix near the nucleosomal DNA entry/exit sites to interact with the Ub at H2AK119 **(Fig. 1d)**.

### Interactions between SSX1 and the nucleosomal H2A/H2B surface

The H2A/H2B acidic patch serves as a hotspot for recruiting many nucleosome effector proteins^24–26^. In our structure, SSX1 is anchored to the nucleosomal H2A/H2B acidic patch by four amino acids (residues W164, R167, L168, R169) **(Fig 2a, Extended Data Fig. 3f, 4a)**. Among them, SSX1 W164 forms a hydrogen bond with H2B Q47 and a hydrophobic interaction with H2A A60. SSX1 R167 forms electrostatic contacts with H2B E113 **(Fig. 2b and Extended Data Fig. 4b)**. SSX1 L168 interacts with the main chain of the H2A α2 helix through van der Waals forces. The SSX1 R169 side chain inserts into an acidic pocket composed of H2A residues E61, D90 and E92, which establishes a salt bridge network **(Fig. 2e and Extended Data Fig. 4c)**. SSX1 mutants at these residues (W164A, R167A, L168A, or R169A) all failed to pull down H2AK119Ub nucleosomes **(Fig. 2c)**, confirming the critical role of these amino acids in mediating nucleosome H2A/H2B acidic patch recognition. The above data also provided the structural basis to support the results of previous photocrosslinking and pull-down experiments on the role of the SSX1 RLR motif (residues R167, L168, R169) in H2AK119Ub nucleosome binding^20^.

**Fig. 2.**
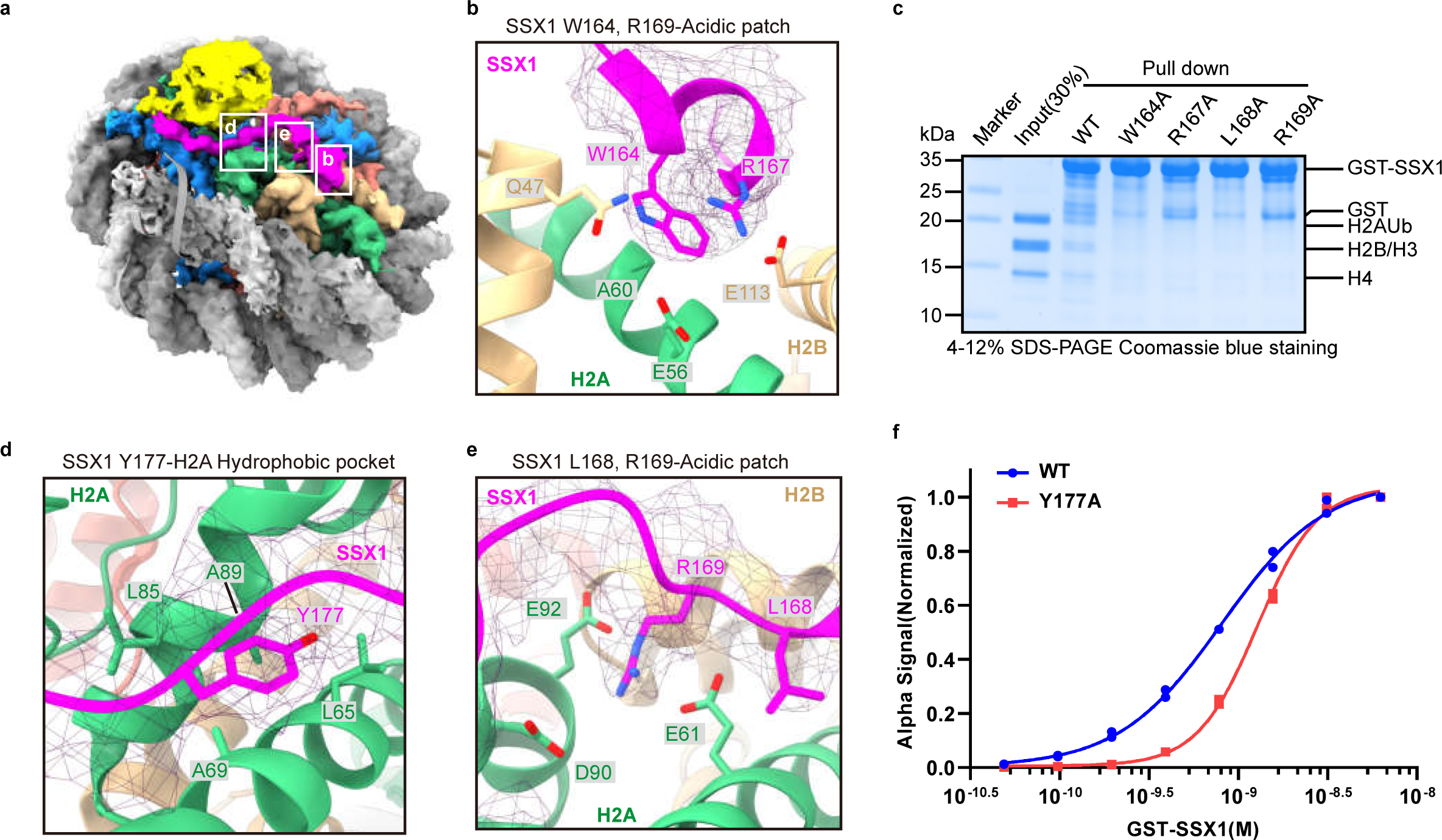
The N-terminal and middle regions of SSX1 interact with the H2A/H2B surface. **a.** Overview of the interaction of SSX1 with the nucleosome surface. White lines frame the three critical interactions region between SSX1 and H2A/H2B surfaces. **b, d, e.** Close-up views of the contacts between SSX1 and the H2A–H2B acidic patch and H2A hydrophobic pocket. **(b)** SSX1 (residues W164, R167) interacts with H2B (residues Q47, E113) and H2A (residues E56, A60); **(d)** SSX1 (residues Y177) interacts with H2A (residues L65, A69, L85, A89); **(e)** SSX1 (residues L168,R169) interacts with the main chain of H2A α2 helix and H2A (residues E61,D90,E92). **c.** Pull-down experiments of WT GST-SSX1 and GST-SSX1 mutants (including W164A, R167A, L168A, R169A) incubated with H2AK119Ub nucleosomes and visualized with Coomassie blue-staining. These gels are representative of three independent experiments. The gel source data is shown in Source Data Fig.1. **f.** AlphaLISA assays performed with GST-SSX1 WT or Y177A mutant and biotinylated H2AK119Ub nucleosomes. Binding curves are plotted as individual data points (n□=□2 assay technical replicates).

Adjacent to the acidic pocket, the side chain of SSX1 residue Y177 was found to insert into a hydrophobic pocket formed by residues L65/A69 of the H2A α2 helix and residues L85/A86 of the H2A α3 helix **(Fig. 2d and Extended Data Fig. 3h, 4d)**. Consistent with this structural observation, AlphaLISA binding assays showed that the SSX1 Y177A mutation reduced the binding affinity to H2AK119Ub nucleosomes **(Fig. 2f, Supplementary Fig. 16)**. The tyrosine interaction with the hydrophobic pocket of the H2A α2/α3 helix may constitute a general binding motif for the recruitment of effector proteins to nucleosomes. Indeed, a highly conserved tyrosine residue (Y137) in the ChEx segment of the single-subunit chromatin remodeler CHD1^27^ and a tyrosine residue (Y1379) in the SMARCA4 SnAc domain of the chromatin-remodeling PBAF complex^28^ also pack into the hydrophobic pocket of the H2A α2/α3 helix **(Extended Data Fig. 4d,e)**.

Collectively, two interactions were observed between SSX1 and the nucleosomal H2A/H2B surface, namely, the interaction of SSX1 N-terminal basic regions (residues W164, R167, L168, R169) with the H2A/H2B acidic patch, and the interaction of SSX1 Y177 with the hydrophobic pocket of the H2A α2/α3 helix. Notably, sequence alignment indicated that the SSX1 residues (W164, R167, W168, R169, Y177) involved in the two interactions are conserved among the homologues SSX2-9 **(Extended Data Fig. 4f)**.

### The H3-Ub basic groove wraps the acidic C-terminus of SSX1

Our structure shows that the C-terminal tail of SSX1 is wrapped by both the Ub at H2AK119 and the histone H3 αN-helix near the nucleosomal DNA entry/exit site **(Fig. 3a, b, Extended Data Fig. 3g)**. Residues R42, R72, and R74 of Ub and residues R49, R52, and K56 of the H3 αN-helix involved in this wrapping interaction are all positively charged, forming a cryptic basic groove that tightly wraps the acidic tail of SSX1 (residues 181-184, SDPE) **(Fig. 3e)**. SSX1 S181 forms a hydrogen bond with H3 K56. SSX1 D182 makes electrostatic contact with H3 R52 and is spatially adjacent to H3 R49. Finally, SSX1 E184 is wrapped by Ub R42, R72, and R74 **(Fig. 3f)**.

**Fig. 3.**
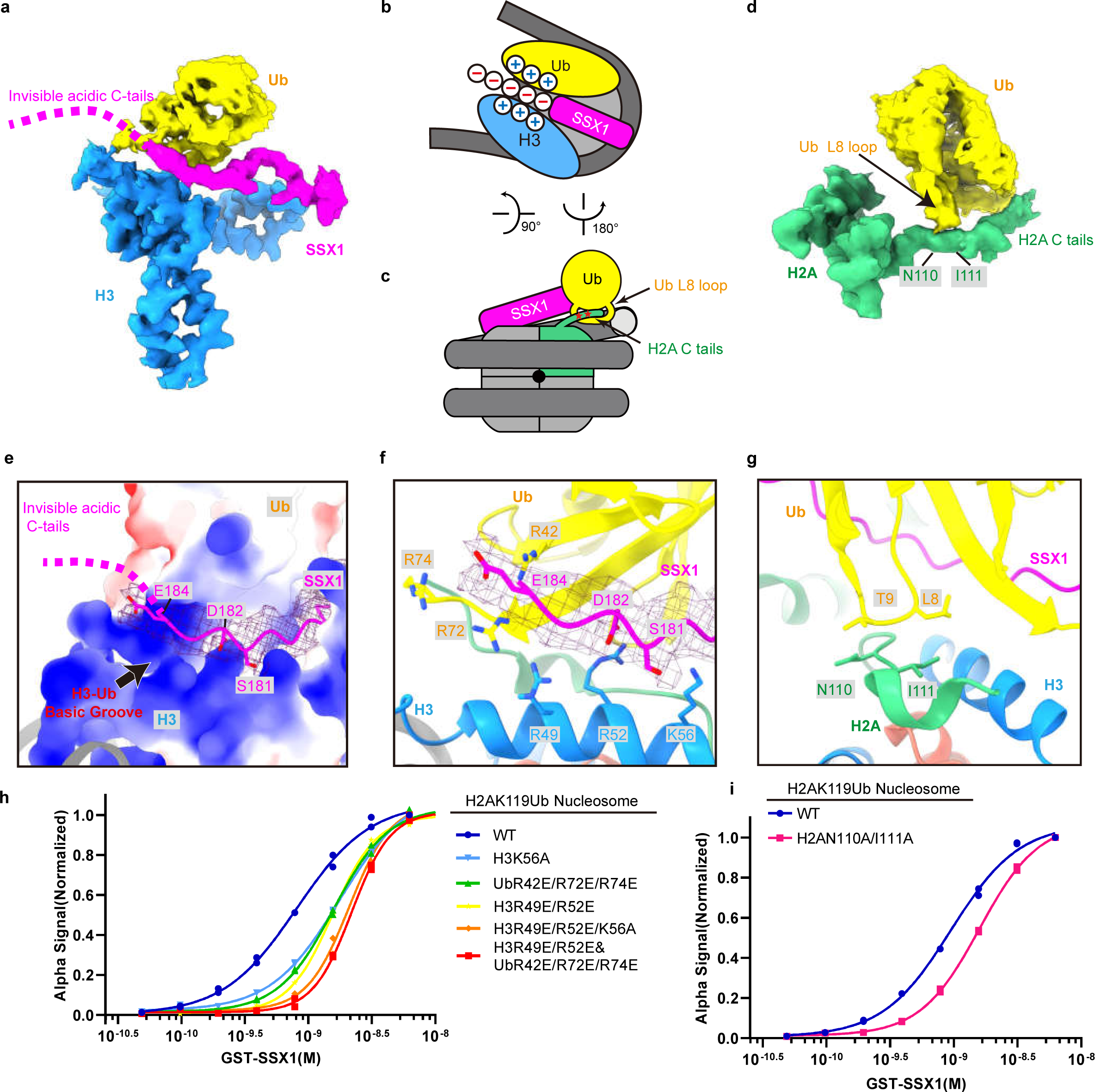
The C-terminal acidic tail of SSX1 interacts with the basic groove formed by histone H3^R49,R52, K56^ and Ub^R42, R72, R74^. **a.** Cryo-EM density map representation of SSX1 interacting with H3-Ub. The density of SSX1 (residues 185-188) is invisible. **b.** Cartoon representation of the SSX1 acidic tail (residues 181-184) wrapped by H3-Ub basic amino acids. **c.** Cartoon representation of the L8 loop of Ub interacting with H2A. **d.** Cryo-EM density map representation of Ub interacting with H2A. **e.** Close-up views of the contacts between SSX1 and the H3–Ub basic groove. The histone octamer and Ub are presented as surface electrostatic potential, positively charged surface coloured in blue and negatively charged surface in red. SSX1 and DNA are depicted in cartoon model. The residues S181, D182, and E184 of SSX1 in stick representation. **f.** Close-up views of the contacts between SSX1 and the H3–Ub basic groove. SSX1, H3 and Ub in cartoon model. Relevant side chains in **f** and **g** are shown as sticks. **g.** Close-up views of the contacts between H2A and the Ub. H2A and Ub are depicted in cartoon model. **h, i.** AlphaLISA assays performed with WT GST-SSX1 and biotinylated H2AK119Ub nucleosomes or mutants. Binding curves are plotted as individual data points (n□=□2 assay technical replicates).

Consistent with the structural observations, H2AK119Ub nucleosomes bearing mutations in the interaction regions, i.e., H3 K56A, H3 R49E/R52E and Ub R42E/R72E/R74E, showed decreased binding affinity to SSX1 **(Fig. 3h**, **Supplementary Fig. 16)**. Combined mutations in these regions, including H3 (R49E/R52E/K56A) and H3(R49E/R52E)-Ub(R42E/R72E/R74E) further reduced the binding affinity of SSX1 to the H2AK119Ub nucleosomes **(Fig. 3h, Supplementary Fig. 16)**. Moreover, a previous study reported that mutation of the C-terminal acidic region of SSX1 to alanines (i.e., residues 182-188 D^182^PEEDDE^188^ → AAAAAAA) relieves the preference of SSX1 for H2AK119Ub nucleosomes over unmodified nucleosomes in AlphaLISA assays, and ChIP-Seq and RNA-Seq results showed that deletion of residues 182-188 of SSX1 affects SS18-SSX-specific BAF complex targeting and its consequent gene expression and proliferation^20^. Taken together, the evidence shows that the basic groove formed by H3 and Ub^H2AK119^ plays a key role in the binding of the SSX1 C-terminal acidic region.

Further support for the importance of the H3-Ub^H2AK119^ basic groove comes from the cryo-EM structure of GST-SSX1 bound to the unmodified nucleosomes, for which 2476 images were collected on a 300 kV Titan Krios microscope and processed with RELION 3.1^23^, and a final map was reconstructed at a resolution of 3.0 Å **(Extended Data Fig. 3d, 5, Table 1)**. Comparing the structures of the SSX1-H2AK119Ub nucleosome complex and SSX1-unmodified nucleosome complex **(Fig. 4a, b)**, we found that the SSX1 N-terminal amino acids (residues W164, R167, L168 and R169) in both structures are anchored in the acidic patch of the nucleosomes **(Fig. 4c)**. However, in the structure of the SSX1-unmodified nucleosome complex, SSX1 density after residue 175 (i.e., residues 176-188) is invisible **(Fig. 4c)**. This observation confirms the need for the H3-Ub^H2AK119^ basic groove to bind the SSX1 C-terminal acidic region. It also indicates that in the absence of Ub at the H2AK119 site, the basic H3 residues R49, R52 and K56 are not sufficient to bind to the SSX1 C-terminal acidic region.

**Fig. 4.**
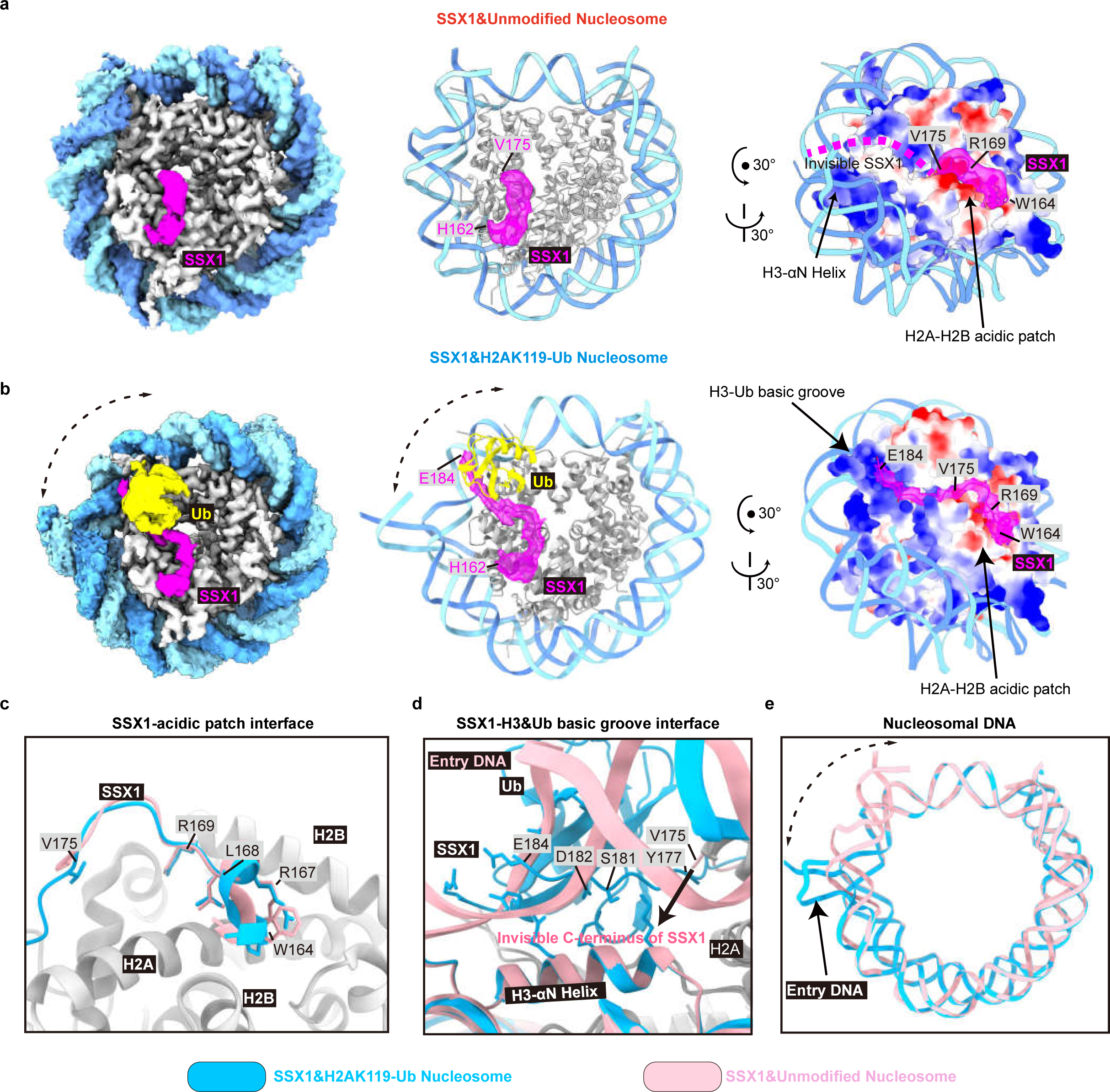
Structural comparison of SSX1-unmodified nucleosome complex and SSX1-H2AK119Ub nucleosome complex. **a.** Structure of SSX1-unmodified nucleosome complex. **(left)** The 3.0 Å Cryo-EM density map of SSX1-unmodified nucleosome complex. **(middle)** Atomic model of the SSX1-unmodified nucleosome complex. The structure is depicted in cartoon with round helices. The density of SSX1 (residues 162-175) is visible. The SSX1 was coloured in magenta, DNA double helix in light sky blue and cornflower blue respectively, and histone octamer in grey. **(right)** Side view of the SSX1-unmodified nucleosome complex. The histone octamer is presented as surface electrostatic potential, positively charged surface coloured in blue and negatively charged surface in red. **b.** Structure of SSX1-H2AK119Ub nucleosome complex. **(left)** The 3.1 Å Cryo-EM density map of SSX1-H2AK119Ub nucleosome complex. **(middle)** Atomic model of the SSX1-H2AK119Ub nucleosome complex. The structure is depicted in cartoon with round helices. The density of SSX1 (residues 162-184) is visible. **(right)** Side view of the SSX1-unmodified nucleosome complex. The density of Ub density is hidden. **c.** Close-up view of comparison of SSX1-acidic patch interface. The structure of the SSX1-unmodified nucleosome complex in **c-e** is coloured in pink. The structure of the SSX1-H2AK119Ub nucleosome complex in **c-e** is coloured in blue. **d.** Close-up view of comparison of SSX1&H3-Ub basic groove interface. **e.** Comparison of nucleosomal DNA in SSX1-unmodified nucleosome complex and SSX1-H2AK119Ub nucleosome complex.

**Table 1.** Cryo-EM data collection, refinement, and validation statistics.

| **Table 1. Cryo-EM data collection, refinement, and validation statistics** |  |  |  |  |
| --- | --- | --- | --- | --- |
|  | SSX1-H2AK119Ub NCP complex (1:1) (EMD-34954, PDB 8HQY) | SSX1-unmodified NCP complex (EMD-34956, PDB 8HR1) | H2AK119Ub NCP (EMD-34957) | SSX1-H2AK119Ub NCP complex (2:1) (EMD-36747) |
| **Data collection and processing** |  |  |  |  |
| Magnification | 81000 | 81000 | 39000 | 81000 |
| Voltage (kV) | 300 | 300 | 200 | 300 |
| Spherical aberration (mm) | 2.7 | 2.7 | 2.7 | 2.7 |
| Detector | K3 | K3 | K2 | K3 |
| Electron exposure (e<sup>-</sup>/Å<sup>2</sup>) | 50.0 e<sup>-</sup>, 32 frames | 50.0 e<sup>-</sup>, 32 frames | 50.0 e<sup>-</sup>, 32 frames | 50.0 e<sup>-</sup>, 32 frames |
| Defocus range (μm) | -1.0 to -2.5 | -1.0 to -2.5 | -1.0 to -1.6 | -1.0 to -2.5 |
| Pixel size (Å) | 1.074 | 1.074 | 0.836 | 1.074 |
| Symmetry imposed | C1 | C1 | C1 | C1 |
| Box size (pixel) | 256 | 256 | 256 | 256 |
| Micrographs (no.) | 4805 | 2476 | 1176 | 4805 |
| Initial particle images (no.) | 3402458 | 2038306 | 165004 | 3402458 |
| Final particle images (no.) | 195091 | 151369 | 23536 | 60624 |
| Map resolution (Å) | 3.1 Å | 3.0 Å | 6.11 Å | 3.6 Å |
| FSC threshold | 0.143 | 0.143 | 0.143 | 0.143 |
| **Refinement** |  |  |  |  |
| Initial model use (PDB code) | 1UBQ,7XD16KW3 | 7XD1 |  |  |
| Model resolution (Å) | 3.24 | 3.15 |  |  |
| FSC threshold | 0.5 | 0.5 |  |  |
| Map sharpening B factor (Å<sup>2</sup>) | -30 | -50 |  |  |
| **Model composition** |  |  |  |  |
| Nonhydrogen atoms | 12303 | 12110 |  |  |
| Protein residues | 848 | 765 |  |  |
| Nucleotides | 272 | 294 |  |  |
| **B factors (Å<sup>2</sup>)** |  |  |  |  |
| Protein | 72.87/180.44/100.11 | 99.08/214.30/120.87 |  |  |
| Nucleotide | 111.63/501.77/175.26 | 134.25/350.98/183.97 |  |  |
| **R.m.s. deviations** |  |  |  |  |
| Bond lengths (Å) | 0.011 | 0.010 |  |  |
| Bond angles (°) | 1.145 | 0.951 |  |  |
| **Validation** |  |  |  |  |
| MolProbity score | 1.61 | 1.64 |  |  |
| Clashscore | 4.42 | 8.44 |  |  |
| Poor rotamers (%) | 0.14 | 0.16 |  |  |
| **Ramachandran plot** |  |  |  |  |
| Favored (%) | 94.2 | 96.92 |  |  |
| Allowed (%) | 5.8 | 3.08 |  |  |
| Disallowed (%) | 0 | 0 |  |  |

Finally, we observed that the L8 loop (residues L8, T9) of Ub forms van der Walls contacts with the αC-helix (residues N110/I111) of histone H2A **(Fig. 3d, g)**. The double mutation of H2A residues N110 and I111 to Ala (H2A N110A/I111A) reduced the binding affinity of SSX1 to the H2AK119Ub nucleosomes **(Fig. 3i, Supplementary Fig. 26)**, suggesting that even a small change in the contacts between Ub and H2A can affect the formation of the H3-Ub^H2AK119^ groove and its ability to bind SSX1. This observation may also explain why SSX1 cannot recognize other ubiquitinated nucleosomes (such as H2BK120Ub, or H2AK127Ub, **Fig. 1b**) because the cryptic H3-Ub^H2AK119^ groove is no longer expected to exist in these systems at all.

### SSX1 mutations affect localization in cells

To perform the cellular validation of our structural studies, we constructed stable 293T cell lines expressing Flag-SS18, Flag-SS18-SSX1(WT), and eight different Flag-SS18-SSX1 mutants. The mutants included variants targeting the nucleosomal acidic patch (W164A, R167A, L168A, R169A), the H2A hydrophobic pocket (Y177A), the H3-Ub basic groove (S181A-D182K-E184R, a deletion of EDDE (residues 185-188) in the S181A-D182K-E184R mutant, and the deletion of seven amino acids (residues 182-188) of SSX1 acidic C-terminal.)

Immunofluorescence (IF) analysis revealed a co-localization between Flag-tagged SS18-SSX1 and H2AK119Ub, while Flag-tagged SS18 exhibited no significant co-localization **(Extended Data Fig. 6a)**. The introduction of mutations in Flag-SS18-SSX1 such as W164A, R167A, L168A, R169A, resulted in a notable reduction or complete loss of co-localization with H2AK119Ub. Additionally, mutations targeting the H2A hydrophobic pocket and the H3-Ub basic groove binding region of SSX1 (Y177A, S181A-D182K-E184R, deletion of EDDE in the S181A-D182K-E184R mutant, deletion of 7 acidic amino acids of SSX1 C terminal) caused almost complete loss of SS18-SSX1 co-localization with H2AK119Ub **(Extended Data Fig. 6a).**

Subsequently, we performed chromatin immunoprecipitation followed by sequencing (ChIP-seq) to obtain genome-wide binding profiles for H2AK119Ub, Flag-SS18, Flag-SS18-SSX1, and eight Flag-SS18-SSX1 mutants. These profiles were then compared to the genomic distribution pattern of H2AK119Ub **(Extended Data Fig. 6b)**. Notably, the binding profiles of wild-type Flag-SS18-SSX1 exhibited enrichment in regions rich in H2AK119Ub. Neither Flag-SS18-SSX1 nor H2AK119Ub displayed a preference for localization at the transcription start site (TSS). However, Flag-SS18 demonstrated a tendency to localize near the TSS. The eight mutants of Flag-SS18-SSX1 exhibited a distinct localization near the TSS similar to Flag-SS18, indicating a clear lack of co-localization with regions abundant in H2AK119Ub. Moreover, Flag-SS18 and Flag-SS18-SSX1 mutants display characteristic retargeting patterns compared to Flag-SS18-SSX1 **(Extended Data Fig. 6c)**.

These findings collectively indicate that the observed mutations in the critical SSX1 interactions with H2AK119Ub nucleosomes have a discernible impact on the cellular localization of SS18-SSX1-containing BAF complex (ssBAF) and its genomic distribution.

### Binding of SSX1 C-terminal tail induces DNA unwrapping

In canonical nucleosomes, the terminal nucleosomal DNA tightly wraps around the histone octamer through contacts with histone residues such as H3 R49 and R52 **(Fig. 5a, Extended Data Fig. 8a)**^29^. However, our SSX1-H2AK119Ub nucleosome complex (1:1 complex) structure shows an exposure of 18 base pairs of DNA from super helix location (SHL) −7.3 to −5.5, causing detachment of the terminal nucleosomal DNA (SHL −7.3 to SHL −5.5) from the pivot of thymine −55, atom P (chain J) by an angle of ∼45° **(Fig. 5c)**. The same behaviour is not observed for the H2AK119Ub nucleosome structure without SSX1 binding (resolved at 6.1 Å, **Supplementary Fig. 18, Table 1**), in which the terminal nucleosomal DNA (SHL −7.3 to SHL −5.5) still tightly wraps around the histone octamer **(Fig. 5b)**. These structural observations suggested that the engagement of SSX1 with H2AK119Ub nucleosomes induces DNA unwrapping. Moreover, in the 2:1 complex, two SSX1 molecules bind to both sides of the nucleosome discs. The two DNA strands at entry and exit sites in the 2:1 complex are observed to be unwrapped **(Extended Data Fig. 7)**. These structural observations suggested that the engagement of SSX1 with H2AK119Ub nucleosomes induces DNA unwrapping.

**Fig. 5.**
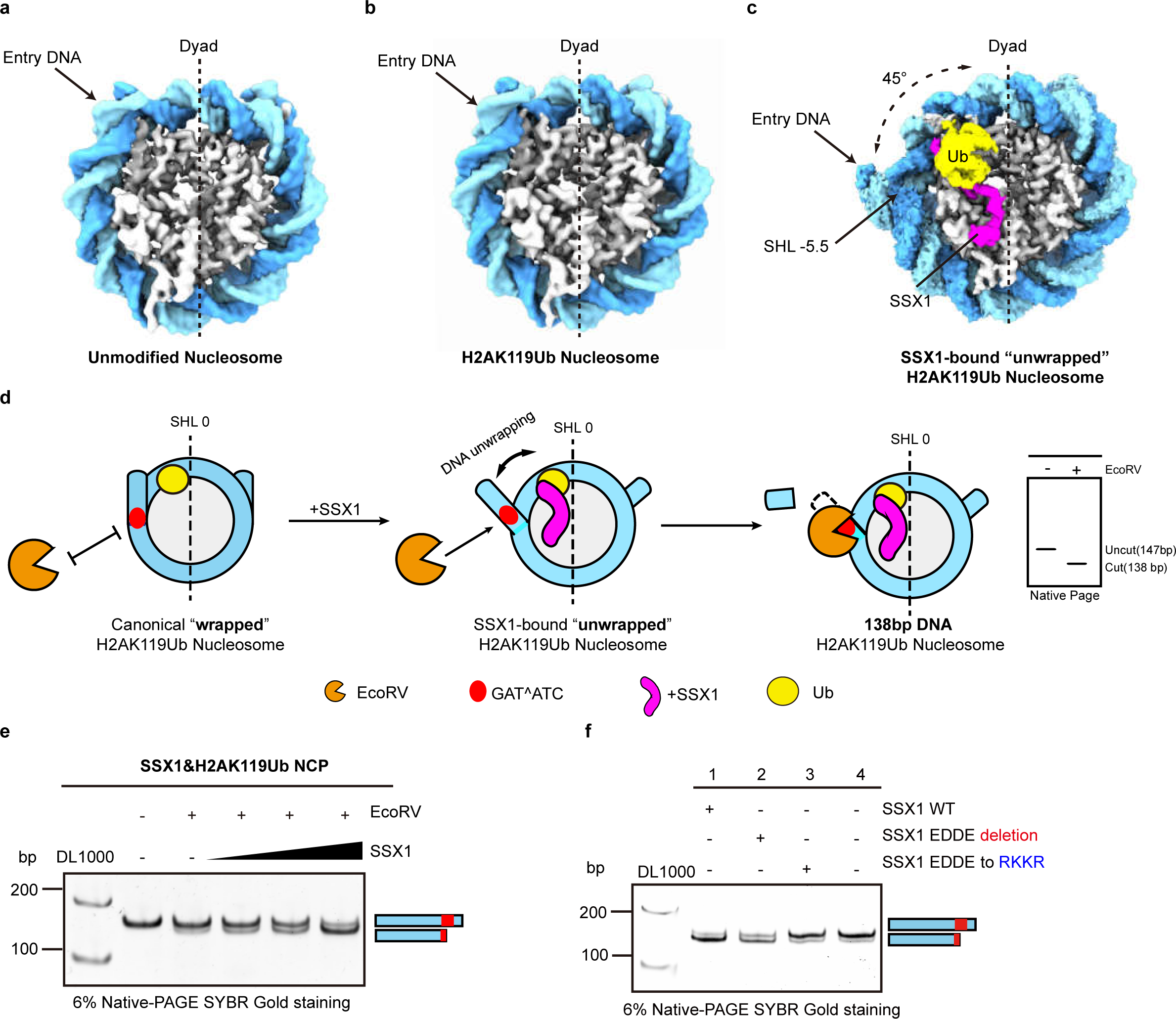
Binding of the C-terminal acidic tail of SSX1 induces nucleosomal DNA unwrapping. **a.** The 3.2 Å Cryo-EM density map of the unmodified nucleosome (EMD-33132). **b.** The 6.1 Å Cryo-EM density map of H2AK119Ub nucleosome. **c.** The 3.1 Å Cryo-EM density map of SSX1-H2AK119Ub nucleosome complex. **d.** Cartoon representation of the schematic of the restriction enzyme accessibility assay (REAA). **(left)** The recognition site is occluded on a canonical “wrapped” H2AK119Ub nucleosome and cannot be cut by EcoRV. **(middle)** The binding of SSX1 to H2AK119Ub nucleosomes induces DNA unwrapping, and the EcoRV recognition site is exposed. **(right)** The nucleosomal 147bp DNA is cut by EcoRV, resulting in 138bp DNA. **e.** SSX1 binding-induced DNA unwrapping was assessed by restriction enzyme accessibility assay (REAA). The schematic of uncut DNA(147bp) and cut DNA(138bp) is shown right. Reactions were analysed by 6% native polyacrylamide gel with SYBR gold staining. This gel is representative of four independent experiments. The gel source data is shown in Source Data Fig.1. **f.** Comparison of DNA unwrapping activity of SSX1, SSX1 (EDDE deletion), and SSX1 (EDDE to RKKR) on 601 H2AK119Ub nucleosomes by REAA. **lane1**: SSX1 (residues 111-188) was incubated with H2AK119Ub nucleosomes, more than 90% of the DNA was cut by EcoRV; **lane2**: SSX1 (residues 111-184) was incubated with H2AK119Ub nucleosomes, approximately 60% of the DNA was cut by EcoRV; **lane3**: SSX1 (residues 185-188, EDDE) to RKKR. was incubated with H2AK119Ub nucleosomes, only 10% of the DNA was cut by EcoRV. **lane4**: H2AK119Ub Nucleosome only. This gel is representative of three independent experiments. The gel source data is shown in Source Data Fig.1.

To validate that SSX1 can induce H2AK119Ub nucleosomal DNA unwrapping, we carried out a restriction enzyme accessibility assay **(Fig. 5d)**^30^. An EcoRV recognition site (GAT|ATC) was introduced into SHL −5.5 of the Widom 601 DNA on the H2AK119Ub nucleosomes. In the absence of SSX1, the GAT|ATC recognition site is occluded in the H2AK119Ub nucleosomes and cannot be cut by EcoRV (< 5% cut after 0.5 hours) **(Fig. 5e)**. Upon the gradual addition of SSX1 to the H2AK119Ub nucleosomes, we observed significant nucleosomal DNA cleavage due to EcoRV digestion, and the cut DNA (138 bp) increased in an SSX1 concentration-dependent manner **(Fig. 5e)**. The biochemical results supported that SSX1 binding to H2AK119Ub nucleosomes induces DNA unwrapping.

In synovial sarcoma cells, SS18-SSX1 localizes to Polycomb-target sites, characterized by a compacted chromatin structure with longer linker DNA and an abundance of linker histone H1. This prompted us to investigate whether SSX1 binding is also capable of triggering DNA unwrapping on 197 bp 25N25 H2AK119Ub nucleosomes and 197 bp 25N25 H1 H2AK119Ub nucleosomes. Encouragingly, we observed an increase of cleaved DNA in an SSX1 concentration-dependent manner on 25N25 H2AK119Ub nucleosomes and 25N25 H1 H2AK119Ub nucleosomes **(Extended Data Fig. 9a-c**). These results suggest that, even within an almost physiologically relevant chromatin context (longer linker DNA and presence of linker histone H1), SSX1 still exhibits the potential to induce nucleosomal DNA unwrapping.

Two factors may contribute to the SSX1-induced terminal DNA unwrapping of the H2AK119Ub nucleosomes. First, in comparing unmodified nucleosomes to 1:1 complex and 2:1 complex, we observed that the side of nucleosome disc with SSX1 binding showed an interaction between H3 (residues R49, R52) and SSX1, while the interaction between H3and terminal nucleosomal DNA (SHL −5.5 to SHL −7.3) was absent **(Extended Data Fig. 8a, b)**. This means that SSX1 competes with terminal DNA for binding to H3, which shall cause the terminal DNA unwrapping. We introduced mutations in H3 residues R49 and R52, replacing them with A or E. Subsequently, we assembled two types of nucleosomes: H3R49A&R52A H2AK119Ub nucleosomes and H3R49E&R52E H2AK119Ub nucleosomes and then performed restriction enzyme accessibility experiments in the absence of SSX1. We found that almost all of the DNA was cut by EcoRV (**Extended Data Fig. 9d**). This result further suggests that the interactions between H3R49&R52 with DNA are necessary for the stable wrapping of the terminal nucleosomal DNA (SHL −5.5 to SHL −7.3).

Second, although not visible in our structure, the negatively charged C-terminus of SSX1 (residues 185-188, EDDE) may electrostatically repel DNA and induce unwrapping. In support of this hypothesis, the SSX1 (111-184) truncation lacking residues 185-188 shows a lower DNA cleavage efficiency on H2AK119Ub nucleosomes than SSX1 (111-188) **(Fig. 5f)**. In addition, almost no cleaved DNA fragments were produced after mutating SSX1 residues 185-188 (EDDE) to RKKR. We hypothesize that it may be that the mutation of EDDE to RKKR in SSX1 has electrostatic interactions with DNA, further reducing DNA unwrapping **(Fig. 5f)**.

## Discussion

Based on our structural and biochemical data, we propose a model describing the mechanism of the selective recognition of H2AK119Ub nucleosomes by SSX1 (**Fig. 6**). First, SSX1 N-terminal basic regions (residues W164, R167, L168, R169) are anchored into the H2A/H2B acidic patch. Second, the aromatic amino acid (residue Y177) in the middle of SSX1 interacts with the hydrophobic pocket formed by H2A α2 and α3 helixes. Finally, and most importantly, the C-terminal acidic region of SSX1 is tightly wrapped by a cryptic basic groove formed by H3 (residues R49, R52, K56) and the Ub motif (residues R42, R72, R74) on the H2AK119 site. This cryptic basic groove constitutes the structural basis by which SSX1 selectively recognizes H2AK119Ub nucleosomes without possessing any Ub-binding domains (**Fig. 6a-c**). This mode of H2AK119Ub recognition is fundamentally different from that previously known for other H2AK119Ub effector proteins that recognize Ub at H2AK119 through their Ub-binding domains (i.e., UIM in JARID2 and zinc finger domain in AEBP2)^10^ **(Extended Data Fig. 10a-c)**.

**Fig. 6.**
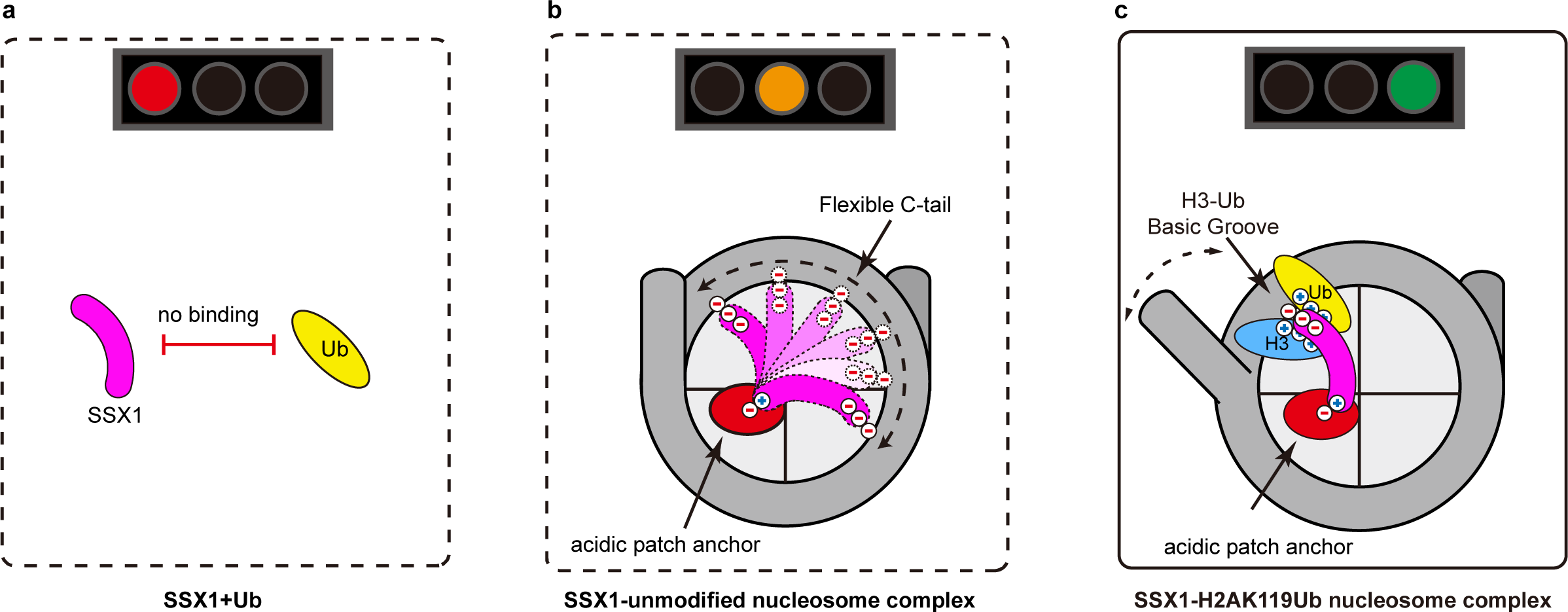
General model of how SSX1 selectively recognizes H2AK119Ub nucleosomes in the absence of the Ub binding domain. **a.** SSX1 is unable to bind free ubiquitin. **b.** On nucleosomes without ubiquitin modification, the N-terminal basic region of SSX1 anchors to the H2A/H2B acidic patch, but the SSX1 C-terminal is flexible and dynamic and cannot stably bind to nucleosomes. **c.** On the H2AK119Ub nucleosomes, in addition to binding the H2A/H2B acidic patch, the C-terminus acidic tails of SSX1 stably bind to the cryptic H3-Ub^H2AK119^ basic groove, allowing SSX1 to recognize H2AK119Ub nucleosomes in the absence of Ub-binding domain.

Comparing the structures of H2AK119Ub nucleosomes in complex with JARID2, AEBP2, and SSX1, we found that Ub motif is situated at fairly different locations **(Extended Data Fig.10d)**. In the structure of JARID2 bound to H2AK119Ub nucleosomes^10^, Ub is located near DNA SHL 0 and just above the H3-αC helix **(Extended Data Fig. 10e)**. However, in the structure of AEBP2 bound to H2AK119Ub nucleosomes^10^, Ub is rotated clockwise by almost 180°close to SHL +1 DNA, and located above the H3-αN helix **(Extended Data Fig. 10e)**. Although Ub in the structure of the SSX1-H2AK119Ub nucleosome complex is also close to SHL −7 DNA and located above the H3-αN helix, Ub in the H2AK119Ub nucleosome complex is much closer to H3 than Ub in the AEBP2 complex (in the SSX1-H2AK119Ub nucleosome complex the distance between H3 residues R49 and Ub R72 is 8.5 Å, while this distance is 22.6 Å in the AEBP2 complex, **Extended Data Fig. 10f**). These observations indicate the plasticity of Ub positions in the complexes of different effector proteins recognizing the same ubiquitinated nucleosomes.

In normal cells, the ATP-dependent chromatin-remodeling BAF complex preferentially resides in the chromatin promoter and enhancer regions and remodels nearby nucleosomes to increase DNA accessibility and facilitate transcription^3,4^. In synovial sarcoma cells, after the fusion of SS18-SSX1, the BAF complex retargets from the enhancers or promoters to the H2AK119Ub-marked polycomb-repressed regions^18^. Previous studies have shown that Ub modification at H2AK119 can reduce the efficiency of terminal DNA unwrapping at DNA entry/exit sites and improve the stability of nucleosomes^31^. However, in our structure, we found that the binding of SSX1 to the H3-Ub^H2AK119^ basic groove induces DNA unwrapping, reshaping the H2AK119Ub nucleosomes from a stable to an unstable conformation, which may in turn facilitate ssBAF remodeling on the H2AK119Ub nucleosomes. Moreover, in our structure, the localization of ubiquitin within the nucleosome is now remarkably precise. This observation may provide valuable insights into the binding model of ssBAF. However, our structural model does not elucidate the effects of SS18-SSX1 incorporation on the global structure of the ssBAF complex and how the ssBAF complex remodels the nucleosome. Therefore, further investigations are required to gain a deeper understanding.

Finally, the H2A/H2B acidic patch and H2A α2/α3 hydrophobic pocket identified in the SSX1-H2AK119Ub nucleosome complex are also employed by other nucleosome interacting proteins (e.g., Dot1L^32,33^, PRC1^34^, and 53BP1^11,35^ for the H2A/H2B acidic patch; SMARCA4^28^ and CHD1^27^ for H2A α2/α3 hydrophobic pocket). It is therefore difficult to find small molecule inhibitors that selectively disrupt the interactions of SSX1 at the interface of the H2A/H2B acidic patch or H2A α2/α3 hydrophobic pocket. In contrast, the SSX1 binding to the H3-Ub^H2AK119^ cryptic pocket is exclusive, making it distinct from other effector proteins. While the H2AK119Ub nucleosome alone fails to stabilize the NCP-Ub ^H2AK119^ interaction, the presence of SSX1 (residues 181-184) induces the formation of the H3-Ub cryptic groove by acting as a molecular glue that binds both Ub and H3^36,37^. The nucleosome-SSX1-Ub triplet interaction is further supported by the interaction between Ub and H2A N110 I111, which helps position Ub appropriately within the H3-Ub cryptic groove. Based on this unique binding mode, unique peptide-like molecular glues can be designed to mimic SSX1(residues 181-184) and competitively bind to the H3-Ub cryptic groove, offering a means for disrupting SSX1’s selective binding to the H2AK119Ub nucleosome. Alternatively, screening for other molecular glues that induce Ub and other histones to interact can also disrupt the H3-Ub cryptic groove, thus providing potential therapeutic options for synovial sarcoma treatment^36^.

## Acknowledgements

We thank the National Key R&D Program of China (No. 2022YFC3401500) for financial support. This study was supported by the National Natural Science Foundation of China (22137005, 92253302, 22227810 for L. Liu, and 22277073 for M. Pan), Shanghai Rising-Star Program (22QA1404900), Shanghai Pilot Program for Basic Research − Shanghai Jiao Tong University (21TQ1400224) and funding from China Postdoctoral Science Foundation (2022TQ0170 and 2022M720075 for H. Ai). We also thank XPLORER prize. H. Ai. thanks the funding by the National Facility for Translational Medicine (Shanghai). We acknowledge the Tsinghua University Branch of China National Center for Protein Sciences (Beijing) for cryo-EM screening and data collection in 200 kV Arctica Tecnai microscopy and 300 kV Titan Krios microscopy. We thank the laboratory of Sheng Ding (School of Pharmaceutical Sciences, Tsinghua University, Beijing 100084, China) for supporting our cellular experiments.

## Author Contributions

1. H. Ai, Z. Tong, M. Pan and L. Liu proposed the idea, designed the experiments and analyzed the results. Z. Tong, H. Ai, Z. Xu and Q. Shi cloned the plasmids, expressed the proteins (SSX1 and histones) and reconstituted the nucleosomes. H. Ai and Z. Tong prepared the cryo-EM samples, collected the cryo-EM data and solved the structures of the H2AK119Ub nucleosomes, SSX1-unmodified nucleosome complex, SSX1-H2AK119Ub nucleosome complex. Z. Tong and Z. Xu performed the pull-downed and AlphaLISA experiments. Z. Tong, H. Ai, and Z. Xu conducted the REAA assay. Z. Xu synthesized the histone H2AK119Ub and H3 mutants. G.-C. Chu synthesized the molecules for CAACU chemical ligation. K. He and Q. Xue conducted cellular experiments (Immunofluorescence and ChIP-seq). Z. Tong and H. Ai, M. Pan and L. Liu wrote and revised the manuscript. All authors read and analyzed the manuscript.

## Methods

### Protein expression and purification

To express the GST-tagged SSX1 and its mutants, DNA sequence coding human SSX1(111-188) or its mutants with codon optimization in *Escherichia coli* species were synthesized and cloned into the pGEX-6P-2 vector (GenScript, Nanjing). The resulting plasmid was transformed into BL21(DE3) competent cells (TransGen Biotech, Beijing), the cells in an LB medium containing 50 μg/mL ampicillin were cultured at 37 □. When the cells were cultured to an OD600 of 1.0, the protein was induced for expression by adding a final 0.4 mM IPTG (isopropylthio-β-galactoside). The cells were cultured at 16 □ and shaken overnight. Cells were harvested by centrifugation and cell pellets were collected and resuspended in GST-lysis buffer (20 mM HEPES, 150 mM NaCl, 1 mM DTT (dithiothreitol), 10% glycerol and 0.1 mM PMSF (phenylmethylsulfonyl fluoride), pH 8.0), sonicated and then centrifugated (2000 x g) to remove the precipitates. The supernatant was incubated with glutathione beads for 2 h at 4 □ and washed with GST-lysis buffer. The target protein was eluted with GST-lysis buffer containing 30 mM glutathione. The elution was diluted with IEX buffer A (25mM HEPES, pH 7.5) to a final 50 mM NaCl, then purified by Mono-Q column-based ion-exchange chromatography (GE Healthcare) (IEX buffer A: 25mM HEPES, pH 7.5; IEX buffer B: 25mM HEPES, 1M NaCl, pH 7.5). The aimed fractions analysed by 15% SDS-PAGE were mixed and concentrated and further purified by size-exclusion chromatography (GE Healthcare) equipped with Superdex 200 Increase 10/300 GL column and pre-equilibrated by SEC buffer (20mM HEPES, pH 7.5, 150mM NaCl, 1mM DTT, 5% glycerol). The peak fractions were collected and stored at −80 □.

For expression of His-tagged SSX1 and its mutants, DNA encoding human SSX1(111-188) or its mutants was cloned into pET28a(+) vector, bearing an N-terminal His×8-tag. Procedure of transformation and cell culture was identical to GST-tagged SSX1 described above, except for the substitution of. Cells were harvested by centrifugation and cell pellets collected were then resuspended in His-tag lysis buffer (20 mM Hepes, 500 mM NaCl, 25 mM imidazole, pH7.5), sonicated and centrifugated (2000 x g). The supernatant was incubated with Ni^2+^-NTA beads (incorporation) for 1 h at 4 □, then washed by His-SSX1 lysis buffer. Bound proteins were eluted by His-SSX1 lysis buffer containing 300 mM imidazole and further purified through Mono-Q-based IEX (IEX buffer A: 25mM HEPES, pH 7.5; IEX buffer B: 25mM HEPES, 1M NaCl, pH 7.5) and Superdex200 Increase column-based SEC (SEC buffer: 20mM HEPES, pH 7.5, 150mM NaCl, 1mM DTT, 5% glycerol) to obtain the target proteins.

Gene sequence (after codon optimization) of wild-type ubiquitin and its mutants (UbG76C) were cloned into pET22b(+) vectors (GenScript, Nanjing). The plasmid was transformed into BL21(DE3) competent cells. The cells were grown in LB medium containing 50 μg/mL ampicillin at 37 □. The protein expression was induced by 1 mM IPTG at 30 □ when the OD600 reached 1.0. The cells were cultured overnight and collected by centrifugation, and the cell pellets were resuspended in ddH_2_O and sonicated for 2 h (100W, 3s on, 5s off). The cell lysate was added with 0.5% v/v HClO_4_ to precipitate impurities. After centrifugation at 2000 x g, the supernatant was dialyzed against 0.1% (v/v) TFA (Trifluoroacetic acid) water solution overnight. The supernatant after dialysis was then purified by Source S 15/150 column-based ion-exchange chromatography (buffer A: 50 mM NaAc, pH 4.5; buffer B: 50 mM NaAc, pH 4.5, 1M NaCl). The eluted fractions containing Ub or its mutant were collected and dialyzed against 50 mM Tris solution (pH 8.0) overnight. The dialyzed product was concentrated and stored at −80 □.

The hUBA1 plasmid (pET3a vector) was transformed into BL21(DE3) cells. The cells were grown at 37 □ in LB media with 50 μg/mL ampicillin. The protein was induced for expression by 0.4 mM IPTG at 18□ overnight. The cell pellets collected by centrifugationwere resuspended in His-tag lysis buffer (20 mM HEPES, 150 mM NaCl, 1 mM DTT, 5% glycerol, 20 mM imidazole, pH 7.5), sonicated and centrifugated (2000 x g). The supernatant was incubated with Co^2+^ beads for 1 h at 4 □, washed with His-tag lysis buffer, then eluted with His-tag lysis buffer containing 300 mM imidazole. The eluent was dialyzed against 20 mM HEPES, 75 mM NaCl, pH 7.5 buffer overnight, then purified by Mono-Q based ion-exchange chromatography (IEX buffer A: 25mM HEPES, pH 7.5; IEX buffer B: 25mM HEPES, 1M NaCl, pH 7.5). Target fractions were mixed and concentrated, stored at −80 □.

The human histones (H2A, H2B, H3 and H4) and their mutants were expressed and purified as previously reported^33,35^.

### DNA preparation

Widom 601 DNA and its variants used to reconstitute nucleosomes were prepared by PCR. Sequences of Widom 601 DNA variants were listed below (the sequence of Widom 601 DNA was underlined).

The sequence of 147 bp Widom 601 DNA: CTGGAGAATCCCGGTGCCGAGGCCGCTCAATTGGTCGTAGACAGCTCTAGCACCGCTTAAACGCACGTACGCGCTGTCCCCCGCGTTTTAACCGCCAAGGGGATTACTCCCTAGTCTCCAGGCACGTGTCAGATATATACATCCTGT

Primer: 5’-CTGGAGAATCCCGGTGCCG-3’

5’-ACAGGATGTATATATCTGACACGTGCC-3’

The sequence of biotinylated Widom 601 DNA: C(biotinylated)CCTGGAGAATCCCGGTGCCGAGGCCGCTCAATTGGTCGTAGACAGCTCTAGCACCGCTTAAACGCACGTACGCGCTGTCCCCCGCGTTTTAACCGCCAAGGGGATTACTCCCTAGTCTCCAGGCACGTGTCAGATATATACATCCTGT

Primer: 5’-C(biotinylated)CCTGGAGAATCCCGGTGCCG-3’ 5’-ACAGGATGTATATATCTGACACGTGCCTGGAG-3’

The sequence of Widom 601 DNA with ECoRV excision site (marked in red): CTGGAGAATCCCGGTGCCGAGGCCGCTCAATTGGTCGTAGACAGCTCTAGCACCGCTTAAACGCACGTACGCGCTGTCCCCCGCGTTTTAACCGCCAAGGGGATTACTCCCTAGT CTCCAGGCACGTGTCAGATA**GAT|ATC**TCCTGT

Primer: 5’-CTGGAGAATCCCGGTGCC-3’

5’-GAACAGGA**GAT|ATC**TATCTGACACGTGCCTGGAGAC-3’

The sequence of 25N25 Widom 601 DNA with ECoRV excision site (marked in red) CTGAGTATCACCCTAGGTCTCTGAT<u>CTGGAGAATCCCGGTGCCGAGGCCGCTCAAT</u>

TGGTCGTAGACAGCTCTAGCACCGCTTAAACGCACGTACGCGCTGTCCCCCGCGTTTTA ACCGCCAAGGGGATTACTCCCTAGTCTCCAGGCACGTGTCAGATA**GAT|ATC**TCCTGTCA CGCGGTGAACAGCGAGATCGGAT

Primer: 5’-CTGAGTATCACCCTAGGTCTCTGATC-3’

5’-ATCCGATCTCGCTGTTCACC-3’

### Histone Octamer reconstitution

The assembly of histone octamer was based on the previous report with minor modifications^38,39^. Briefly, histone H2A (or H2AUb), H2B, H3 and H4 (15 μM each) were weighted and dissolved in histone unfolding buffer (6 M Gn·HCl, 20 mM Tris, 1 mM DTT, pH 7.5), and dialyzed against histone refolding buffer (2 M NaCl, 10 mM Tris, 1 mM EDTA, pH 7.5) at 4 □. The refolding buffer was exchanged with a fresh one every 12 hours. Histone octamer was purified by Superdex 200 Increase 10/300 GL column-based SEC (GE Healthcare) that was pre-equilibrated with refolding buffer. Peak fractions were resolved on 8–16% SurePAGE Bis-Tris (GenScript) gels.

### Nucleosome reconstitution

The procedure of nucleosome reconstitution was consistent with previous work^38,39^, except for minor modifications. Generally, histone octamer and nucleosomal DNA were mixed at a stoichiometric ratio of 1:1 under high salt conditions (approximately 2 M), then the mixture was dialyzed against histone refolding buffer at 4 □ using a dialysis tube (D-Tube^TM^ Dialyzer Medi or Maxi, MWCO 6-8 kDa, Merck). HE buffer (10 mM HEPES, 1 mM EDTA, pH 7.5) was transported to the dialysis vessel at a speed of 0.6 mL/min by a peristaltic pump to gradually decrease the salt concentration. When the salt concentration reached 300 mM or lower, the nucleosomes were purified by ion-exchange equipped with TSKgel DEAE-5PW HPLC column. Peak fractions eluted at around 30 mS/cm conductance were collected and dialyzed against 10 mM HEPES (pH 7.5) overnight. Correct assembly of nucleosome was confirmed by 4.5% native-PAGE with SYBR Gold staining.

### Restriction enzyme accessibility assay

The unmodified or H2AK119Ub nucleosomes (unmodified nucleosome, H2AK119Ub nucleosome, or H3 R49/52A H2AK119Ub nucleosome, H3 R49/52E H2AK119Ub nucleosome,) (500 nM) and GST or His-tagged SSX1 (or its mutant) (3 μM) diluted by 1× CutSmart buffer (New England BioLab) were incubated on ice for 15 mins, then ECoRV restriction enzyme (0.8 U/mL, New England BioLab) was added and incubated at 37 □ for 30 mins. An equal volume of quenching buffer (20 mM HEPES, 4 mM Tris, 10% glycerol, 2% SDS, 10 mM EDTA, pH 7.5) was added to quench the digestion. The mixture was then supplemented by Proteinase K (0.67 mg/mL) and incubated at 37 □ for 30 mins. Finally, the DNA was resolved by 6% native-PAGE and stained by SYBR Gold.

### AlphaLISA binding assay

10 μL of 200 nM (or serially diluted) GST-tagged protein (or its mutants) and 10 μL of 12 nM biotinylated nucleosome (or its mutants) were mixed and incubated in OptiPlate-384 microplate (PerkinElmer) for 60 mins, then 10 μL of 20 μg/mL glutathione acceptor beads (PerkinElmer, AL109C, Lot: 2851471) and 10 μL 20 μg/mL streptavidin donor beads (PerkinElmer, 676002S, Lot: 2838642) were added and incubated for another 1 h. Emission at 615nm was detected by Envision plate reader (PerkinElmer). Two biological replicates were conducted for each binding test. The data was processed in GraphPad Prism-9.0.

### GST-affinity pull-down assay

8 μg of purified GST-tagged SSX1 or its mutants were incubated with 10 μLof glutathione beads (Smart-Lifescience, Changzhou, China) in pull-down buffer (50 mM Tris, 150 mM NaCl, 1 mM DTT, 0.1% (v/v) NP-40, 0.1% (w/v) BSA, pH 7.5) for 30 mins at 4 □, followed with washing with pull-down buffer for 5 times to remove unbound proteins. The SSX1-coated beads were then incubated with 6 μg of ubiquitinated nucleosomes for 2 h at 4 □, and washed with pull-down buffer 4 times. 4×LDS loading buffer containing 100 mM DTT was used to elute the protein with boiling at 95 □ for 5 mins. Histones in pull-down were visualized under Coomassie blue staining.

### Cryo-EM sample preparation

For the SSX1-unmodified nucleosome complex and the SSX1-H2AK119Ub nucleosome complex, we investigated the impact of different SSX: nucleosome ratios (e.g., 1:2, 1:4, 1:6), different nucleosome concentrations (e.g., 4 μM, 2 μM, 0.5 μM, etc.), and different methods of complex incubation (e.g., one-pot addition, step-wise addition) on the acquisition of the sample during cryo-EM sample preparation. Our findings revealed that step-wise addition of SSX1, reaching a final concentration of 0.5 μM nucleosomes with a 6-fold excess, greatly facilitated the formation of complexes, minimizing the presence of individual nucleosomes after glutaraldehyde cross-linking. The gradual step-wise addition method promotes the formation of the complexes better than the one-pot addition. Specifically,6-fold excess of SSX1 was gradationally added into the nucleosome, the SSX1 was added every 10 mins for a total of 5 times. Then the complex was crosslinked with glutaraldehyde (final concentration 0.1% v/v) at 25 □ for 10 mins, the crosslinking reaction was quenched by a final 100 mM Tris (pH 7.5). The crosslinked samples were purified through size-exclusion chromatography by Superose 6 Increase 10/300 GL column that was pre-equilibrated with 20 mM HEPES, 25 mM NaCl, pH 7.5 buffer. Fractions were analysed by 4%-12% SDS-PAGE (Thermo Fischer Scientific) and 4.5% native-PAGE, and the complex-containing fractions were collected and concentrated to 2 μM for further grid preparation. The H2AK119Ub nucleosome sample was prepared similarly.

The cryo-EM sample was prepared using the Vitrobot system (Thermo Fischer Scientific). 3.5 μL sample was applied to a glow-discharged 300 mesh gold grid (Quantifoil R1.2/1.3). The sample was incubated with the grid for 60 seconds at 8 □ and 100% humidity to maximize the absorption. Then the grids were plunged into liquid ethane and stored in liquid nitrogen.

### Data collection and processing

4805 images for the SSX1-H2AK119Ub nucleosome complex and 2476 images for the SSX1-unmodified nucleosome complex were collected on a 300 kV Titan Krios electron microscope (FEI Company) equipped with a Gatan K3 direct electron detector at a nominal 81,000× magnification. The calibrated pixel size was 1.074 Å. 1176 images for H2AK119Ub nucleosome were collected on a 200 kV Tecnai Arctica microscope equipped with a K2 camera, the pixel size was 0.836 Å. The image was recorded for 32 frames in a total of 2.56 seconds. The defocus was set between −1.0 and −2.5 μm. The beam-induced motion was timely processed with MotionCor2 during data collection. The cryo-EM datasets were all processed with RELION 3.1^23^. Kai Zhang’s Gctf-1.18^40^ was used to estimate the CTF parameters. The C1 symmetry was used for data processing. For each dataset, manual picking was employed to give the 2D reference for particle auto-picking. The particles after two rounds of 2D classification using fast subsets in RELION were used for 3D initial model generation, which was further applied for 3D classification. The binned particles were processed to give the desired conformation with multi-rounds of 3D classifications. The final good-quality particles were re-extracted to bin1 to reconstitute the final high-resolution map. The cryo-EM processing flowcharts of the SSX1-H2AK119Ub nucleosome complex (1:1 complex and 2:1 complex), the SSX1-unmodified nucleosome complex and the H2AK119Ub nucleosomes were shown in **Extended Data Fig. 2, 5 and Supplementary Fig. 18**. The resolution was reported with the FSC 0.143 criteria. ResMap-1.14 was used to estimate the local resolution of the cryo-EM maps. The cryo-EM data processing statistics were summarized in **Table 1**.

### Model building and refinement

To build the atomic model of SSX1-H2AK119Ub nucleosome complex and SSX1-unmodified nucleosome complex, the nucleosome structure (PDB: 7XD1, 6KW3) and/or Ub structure (PDB: 1UBQ) were rigid-body docked in the cryo-EM map in ChimeraX-1.2.5^41^. The coordinates were exported relative to the cryo-EM maps to reserve the position information. Then the nucleosome and Ub coordinates were opened in WinCoot-0.8.2^42^ and merged into one molecule. On this basis, the SSX1(162-184) was manually built according to the main chain traveling direction and sidechain orientations. Especially, the residue Y177 shows a representative amino acid with its aromatic side chain inserted into the H2A α2/α3 helix. Then the rough model was manually inspected and refined with Phenix-1.13^43^ using real-space refinement. The final models of SSX1-H2AK119Ub nucleosome complex or SSX1-unmodified nucleosome complex were validated in Phenix-1.19.2^43^. The cryo-EM map and model were viewed in ChimeraX-1.2.5^41^, in which the images were exported for painting.

### Plasmid construction

All plasmids were constructed with NEBuilder® HiFi DNA Assembly Master Mix (New England BioLabs) according to the manufacturer’s instructions. Briefly, DNA fragments with about 20bp overlapping ends were generated by restriction enzyme digestion or PCR. Then DNA fragments were mixed with the Assembly Master Mix in a volume ratio of 1:1 and incubated at 50°C for approximately 30 minutes followed by transforming Trans1-T1 Phage Resistant Chemically Competent Cell (TransGen Biotech) and Sanger sequencing.

### Lentivirus production

Lentiviral vectors were transfected together with helper plasmids pMD2.G (#12259) and psPAX2(#12260) into HEK293T cells to produce lentivirus. 36 hours later, supernatant was collected and filtered through a 0.45-μm filter. Lentiviral were stored at −80 °C or used for stable cell line construction.

### Cell culture and stable cell line construction

HEK293T cells (ATCC, Lot: CRL-11268) were cultured in DMEM (Invitrogen) supplemented with 10% FBS (Invitrogen), and 1% penicillin/streptomycin (Invitrogen) at 37°C with 5% CO2 incubation. To generate stable cell line, HEK293T cells were infected with lentivirus. 24 hours after infection, cells were selected by 1 ug/ml Puromycin (Selleck) for 2days and maintained at 0.5 ug/ml Puromycin for 7 days.

### Chromatin IP-seq

The chromatin immunoprecipitation (ChIP) assay was performed with SimpleChIP® Plus Enzymatic Chromatin IP Kit (Magnetic Beads) (Cell signaling technology, #9005) following the manufacturer’s instructions. Briefly, approximately 4×10^6^ cells (a 15 cm culture dish) were used for each ChIP assay. Cells were fixed with 1% formaldehyde for 10 min at room temperature. After quenching by adding 2 ml of 10X glycine for 5 min at room temperature, cells were scraped, lysed, digested and sheared by sonication. Clarify lysates by centrifugation, transfer 2% supernatant as Input Sample, which can be stored at −20°C until further use. Lysates supernatant was incubated overnight at +4□°C with immunoprecipitating antibody (Ubiquityl-Histone H2A (Lys119) (D27C4) XP® Rabbit mAb #8240, DYKDDDDK Tag (D6W5B) Rabbit mAb #14793 (Cell Signalling technology) diluted at 1:100 and 1:50 ratios, respectively) followed by adding 30 µl of Protein G Magnetic Beads to each IP reaction and incubating for 2h at 4°C with rotation. After washing the beads, chromatin was eluted from the antibody/protein G magnetic beads. To all samples, including the 2% input sample from previous steps, reverse cross-links by adding 6 µl 5M NaCl and 2 µl Proteinase K, and incubate 2 h at 65°C followed by DNA purification using Spin Columns. These immuno-enriched DNA samples could be used for NGS after DNA library construction with NEBNext® UltraTM II DNA Library Prep Kit (NEB, USA, Catalog #: E7645L). The qualified libraries were pooled and sequenced on Illumina platforms with PE150 strategy in Novogene Bioinformatics Technology Co., Ltd (Beijing, China). The raw reads were performed quality control using Fastp (0.20.1), then aligned to hg19 genome by hisat2 (2.2.1). Peak calling was performed with macs2 (2.2.7.1) and deeptools (3.5.1)..

### Immunofluorescence

Cells were washed with PBS, fixed with 4% paraformaldehyde at room temperature for 15 min, permeabilized and blocked with 0.2% Triton X-100 and 3% BSA in PBS for 1h. After washing 3 times with PBS, cells were incubated with primary antibody (Ubiquityl-Histone H2A (Lys119) (D27C4) XP® Rabbit mAb #8240, DYKDDDDK Tag (D6W5B) Rabbit mAb #14793) (Cell Signalling technology)diluted in the blocking buffer at 1:1000 radios for 2 hours. Cells were washed with PBS 3 times and incubated with secondary antibodies (Donkey anti Mouse-488 #A-21202, Donkey anti Rabbit-555 #A-31572) (Thermo Fischer Scientific) diluted in the blocking buffer at 1:1000 radios for 1h, then incubated with 1ug/ml DAPI for 5min. After washing with PBS 3 times, cells were performed fluorescence microscopy.

## Data availability

Cryo-EM maps have been deposited in the Electron Microscopy Data Bank (EMDB, www.ebi.ac.uk/pdbe/emdb/) under the accession codes EMD-34954 (SSX1-H2AK119Ub NCP complex, 1:1), EMD-34956 (SSX1-unmodified NCP complex), EMD-34957 (H2AK119Ub NCP), EMD-36747 (SX1-H2AK119Ub NCP complex, 2:1). The atomic models have been deposited in the Protein Data Bank (PDB, www.rcsb.org) under the accession codes 8HQY (SSX1-H2AK119Ub NCP complex), 8HR1 (SSX1-unmodified NCP complex).

All ChIP-seq data have been deposited in the the GEO repository under the GEO accession number: GSE236811 (https://www.ncbi.nlm.nih.gov/geo/query/acc.cgi?acc=GSE236811).

---

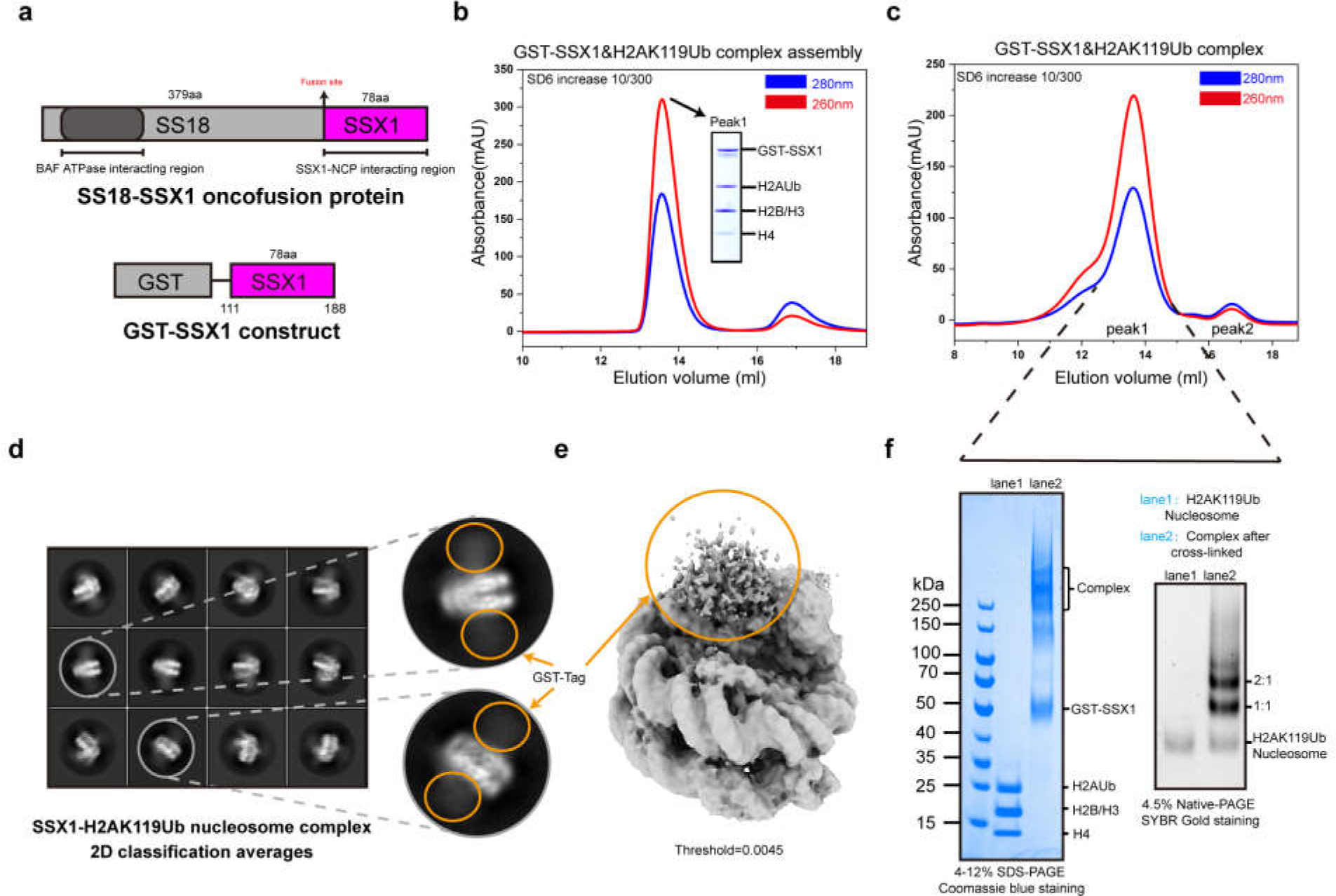

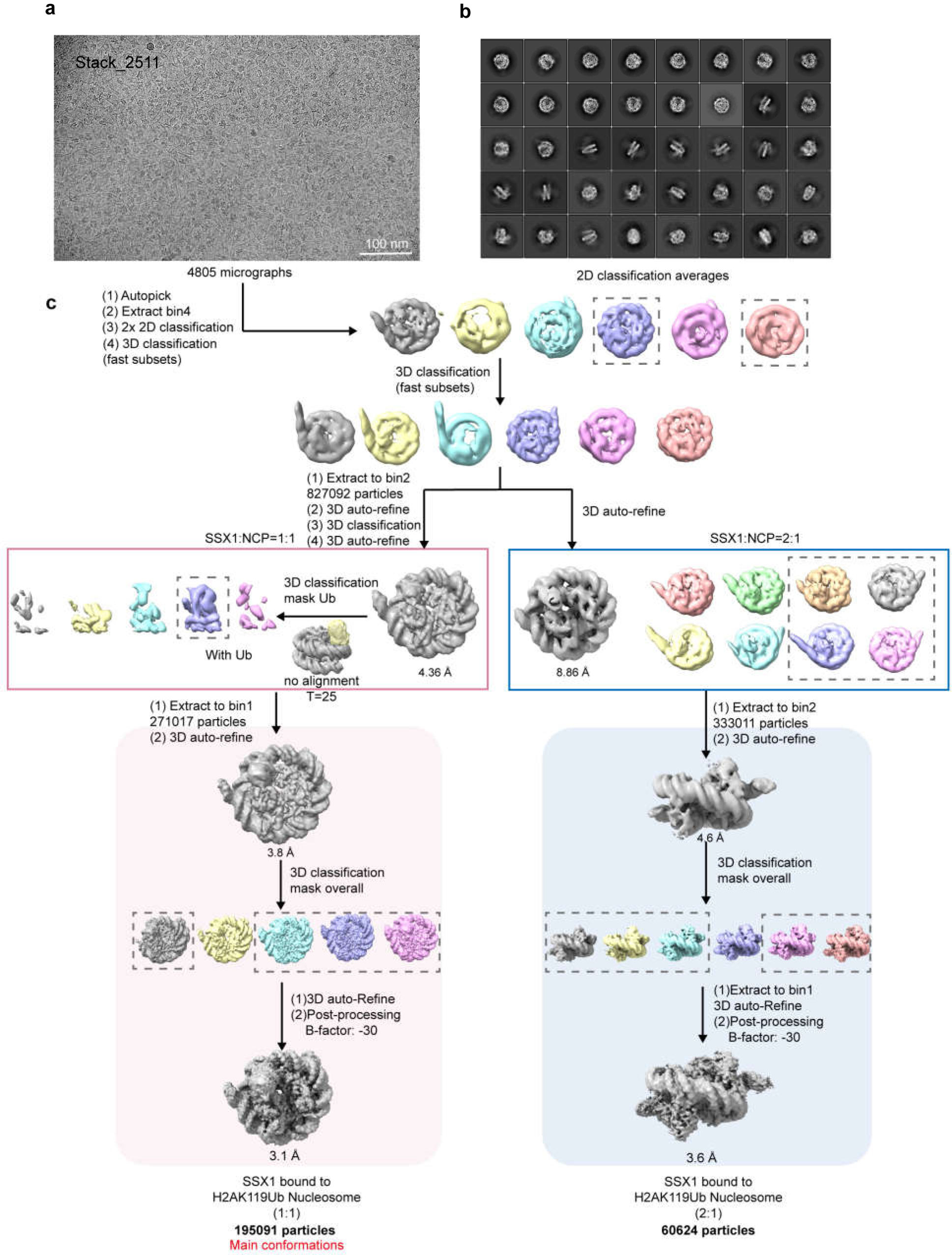

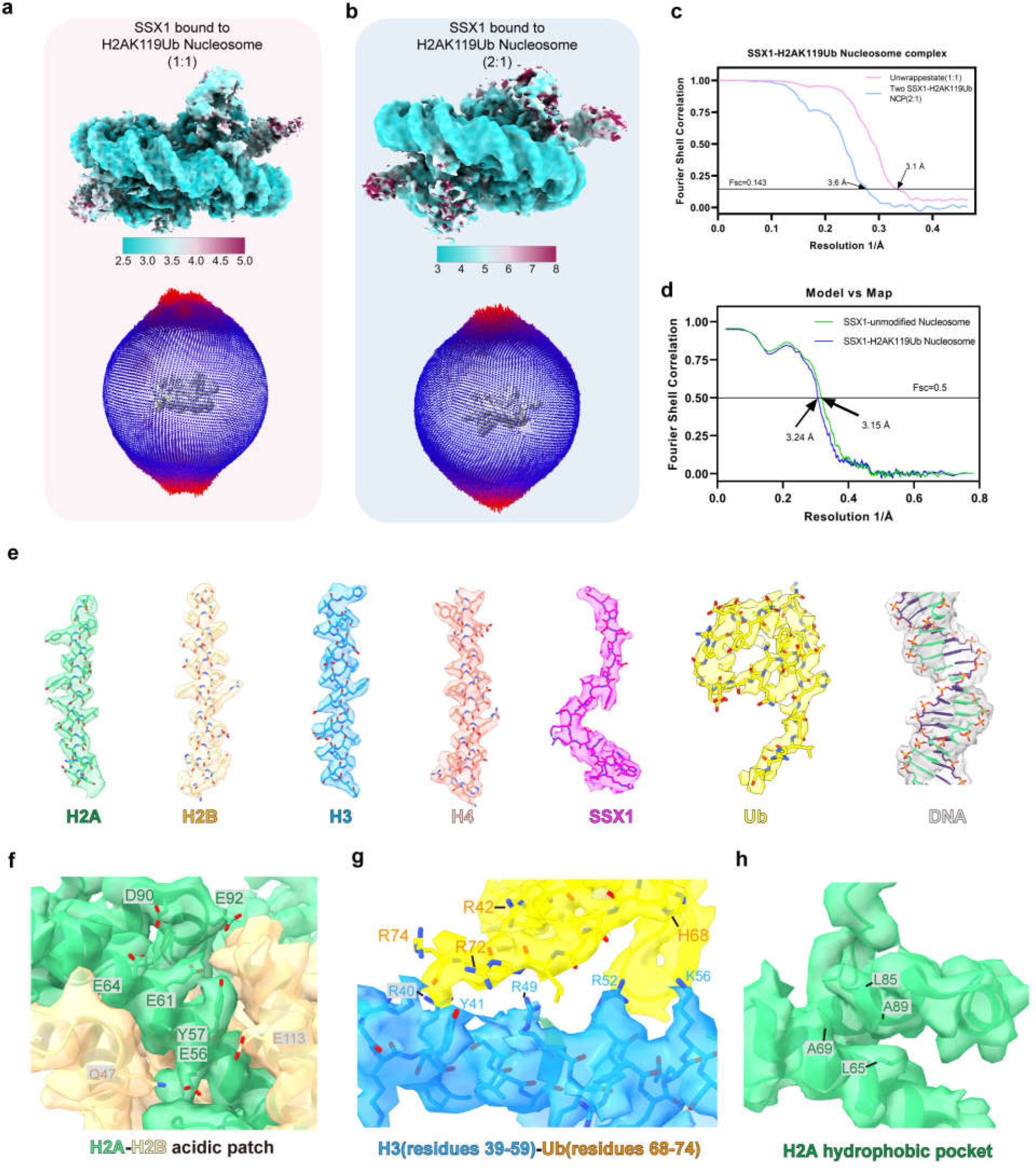

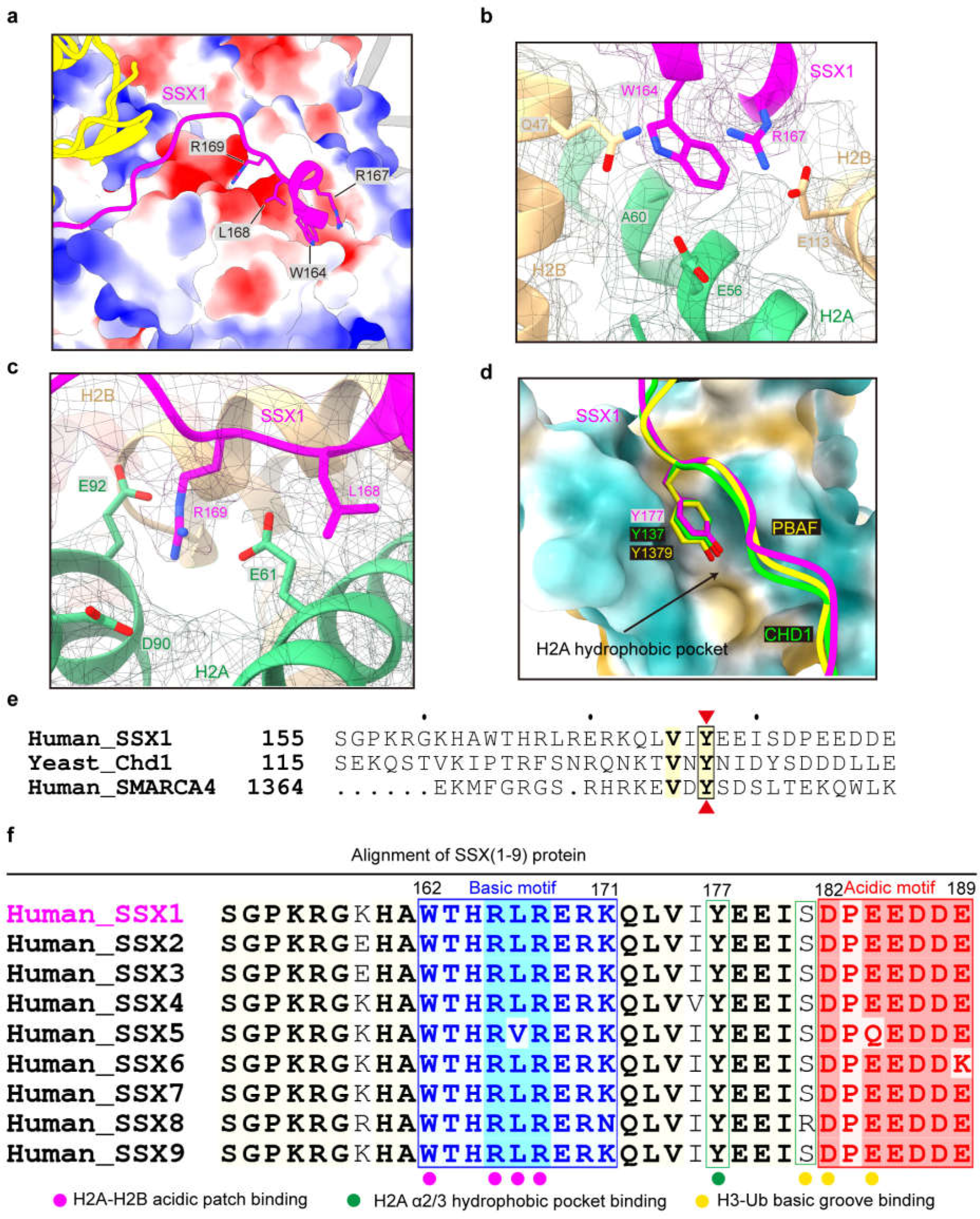

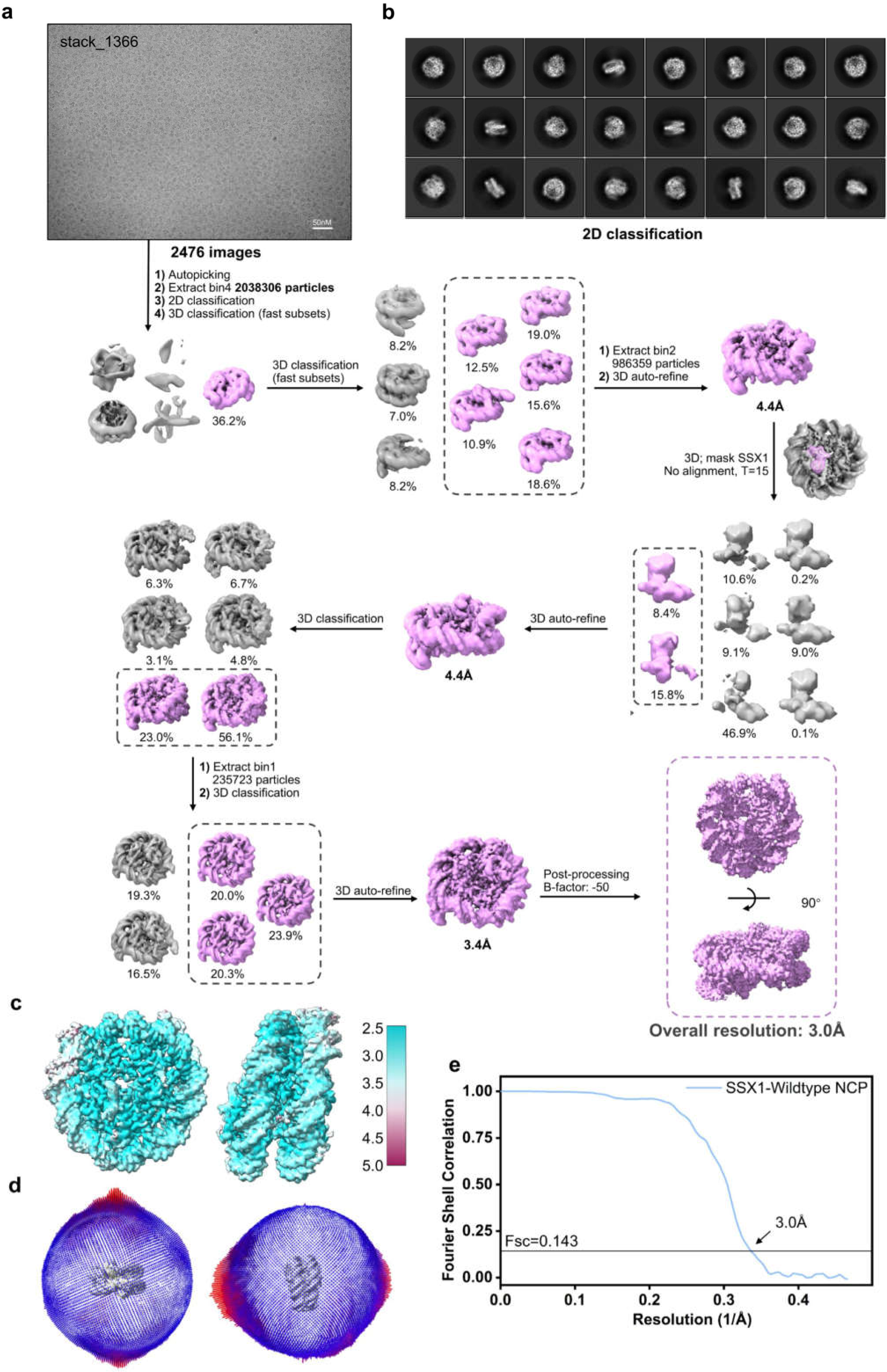

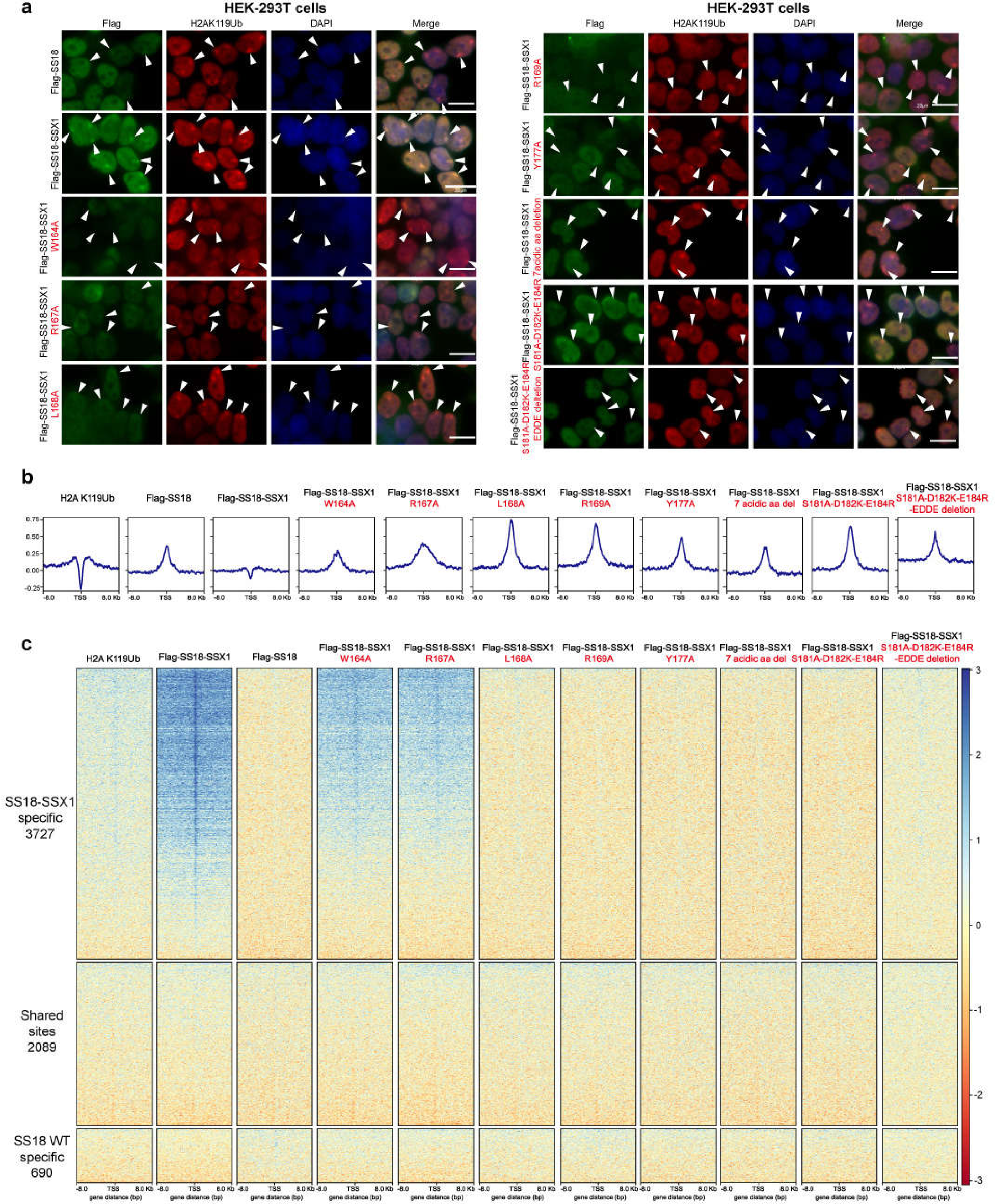

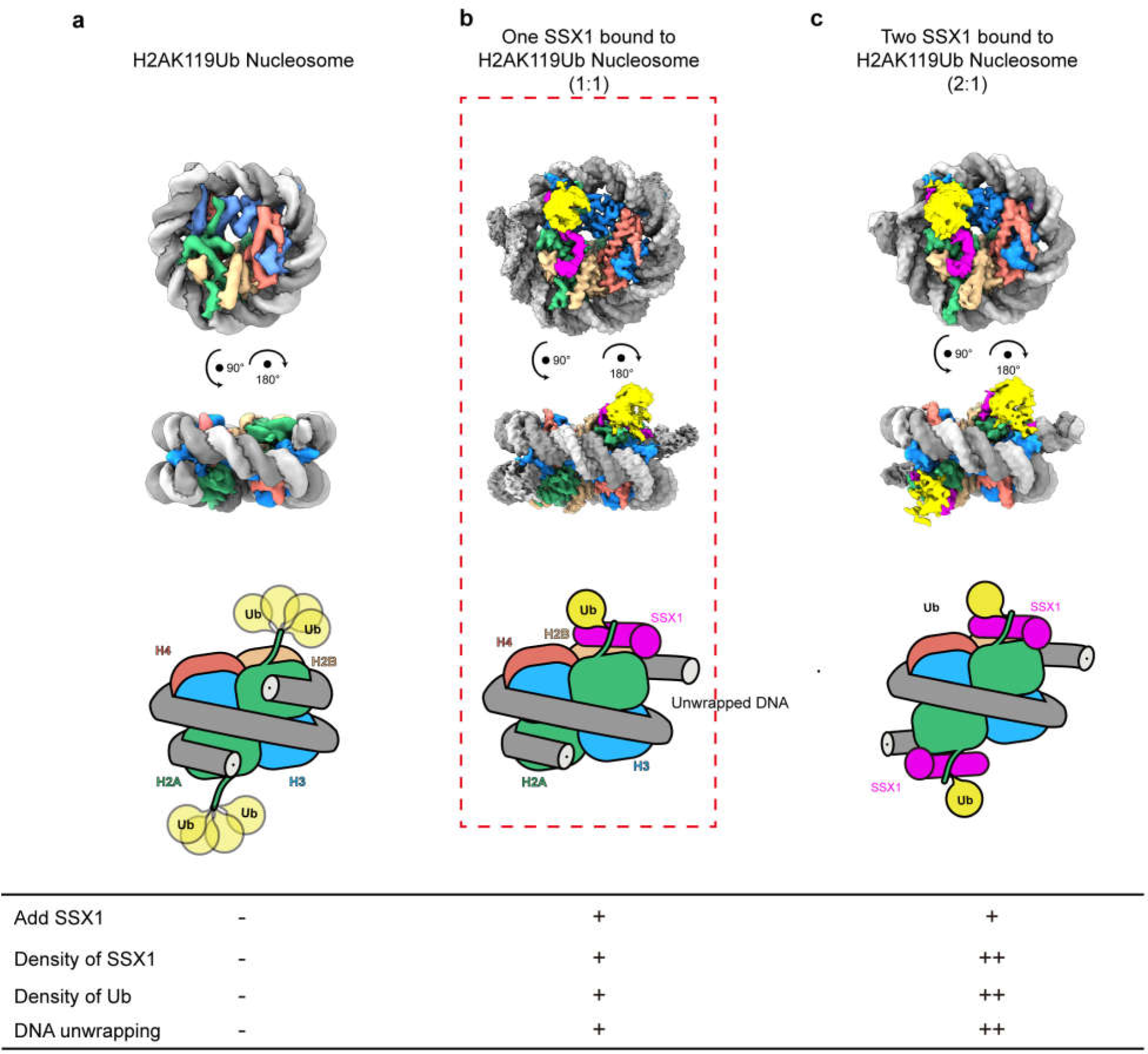

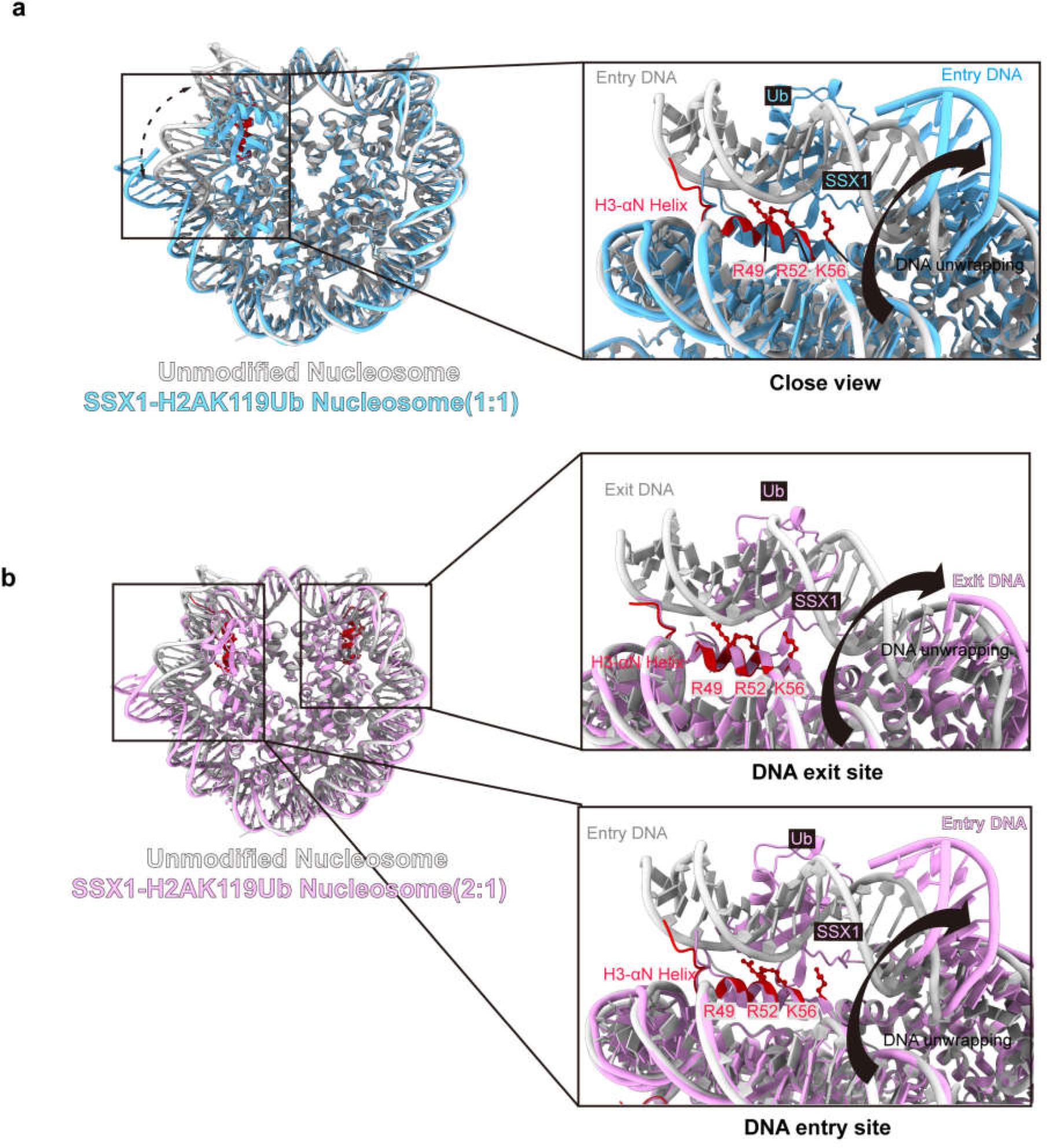

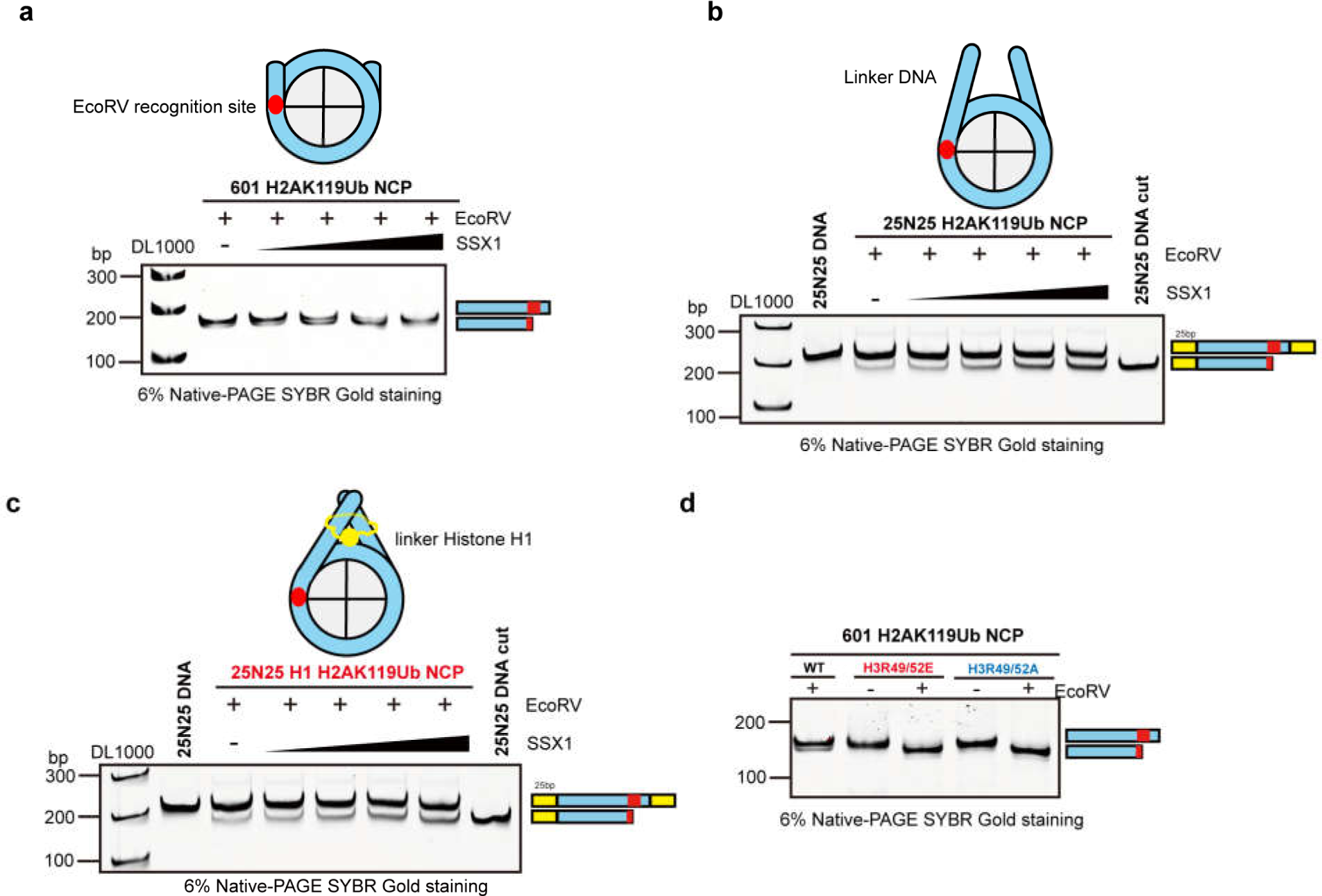

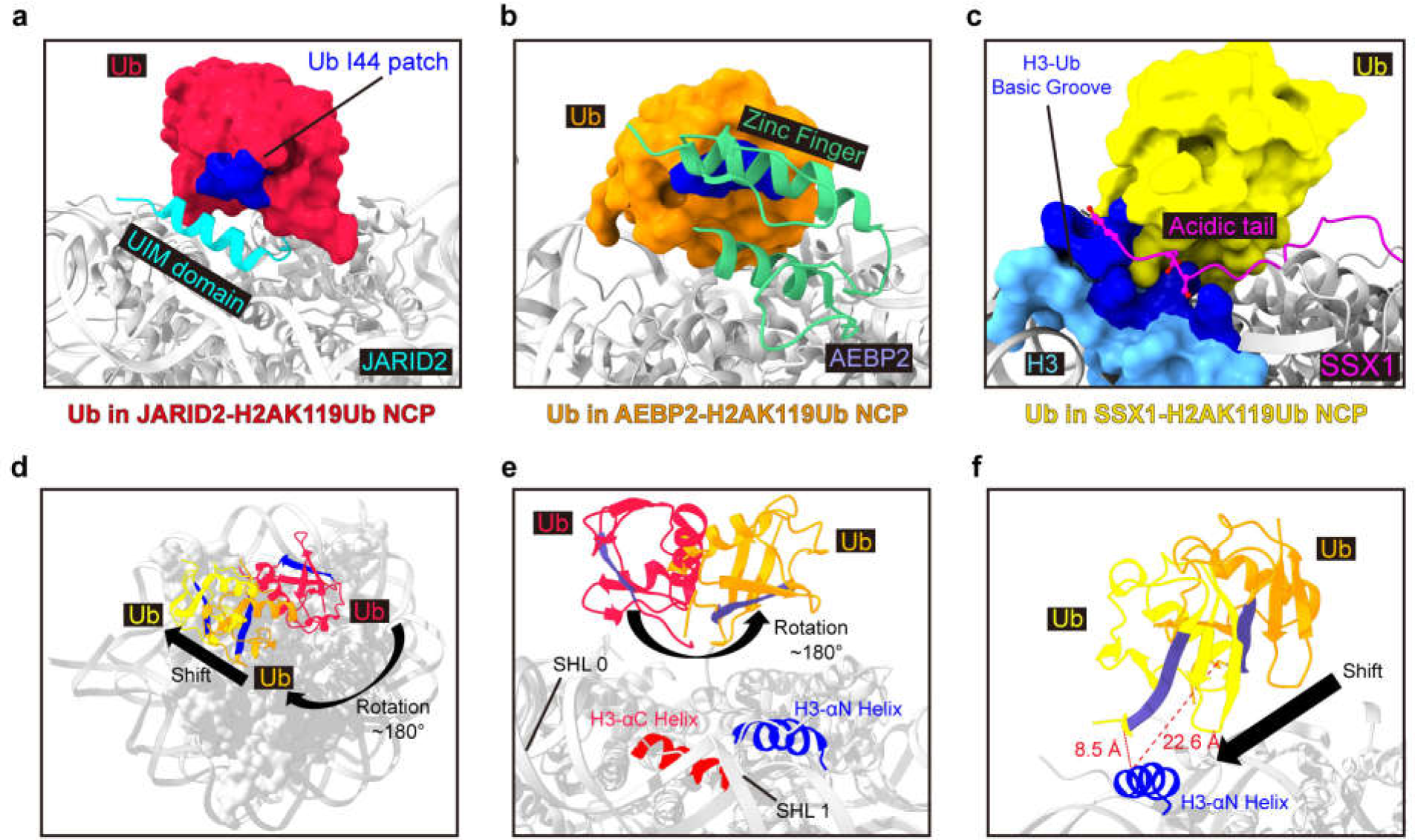

